# Minimization of gene editing off-target effects by tissue restriction of expression

**DOI:** 10.1101/2025.01.21.634017

**Authors:** Nam-Gyun Kim, Martine Aubert, Anoria K. Haick, Paola A. Massa, Michelle A. Loprieno, Marius Walter, Hyeong Geon Kim, B. Ethan Nunley, Hong Xie, Laurence Stensland, Lindsay M. Klouser, Ailyn C. Pérez-Osorio, Daniel Stone, Pavitra Roychoudhury, Keith R. Jerome

## Abstract

Therapeutic *in vivo* gene editing with highly specific nucleases has the potential to revolutionize treatment for a wide range of human diseases, including genetic disorders and latent viral infections like herpes simplex virus (HSV). However, challenges regarding specificity, efficiency, delivery, and safety must be addressed before its clinical application. A key concern is the risk of off-target effects, which can cause unintended and potentially harmful genetic changes. We previously developed a curative *in vivo* gene editing approach to eliminate latent HSV using HSV-specific meganuclease delivered by an AAV vector. In this study, we investigate off-target effects of meganuclease by identifying potential off-target sites through GUIDE-tag analysis and assessing genetic alterations using amplicon deep sequencing in tissues from meganuclease treated mice. Our results show that meganuclease expression driven by a ubiquitous promoter leads to high off-target gene editing in the mouse liver, a non-relevant target tissue. However, restricting the meganuclease expression with a neuron-specific promoter and/or a liver-specific miRNA target sequence efficiently reduces off-target effects in both liver and trigeminal ganglia. These findings suggest that incorporation of regulatory DNA elements for tissue-specific expression in viral vectors can reduce off-target effects and improve the safety of therapeutic *in vivo* gene editing.

## INTRODUCTION

Gene editing technologies, mediated by highly specific nucleases such as CRISPR-family proteins, meganucleases, zinc finger nucleases, or TAL effector nucleases, offer unprecedented potential to treat a wide range of human diseases by precisely modifying the genetic code.^1–3^ *Ex vivo* gene editing is being increasingly used to engineer immune cells to target cancer, or to correct genetic defects in hematopoietic or other stem cells for transplantation.^4–9^ However, some of the most transformative applications of gene editing would require it to occur *in vivo*, allowing the direct modification of genes within living organisms.^4–6^ Recent advancements in delivery methods, such as viral vectors and nanoparticle systems, have significantly improved the efficiency and specificity of *in vivo* gene editing.^6,10,11^ However, significant challenges remain to be addressed before *in vivo* gene editing can reach its full promise.^4–6,12^

One of the most important concerns in gene editing is the possibility of off-target effects, due to their potential to cause unintended genetic alterations.^4–6^ These effects occur when the gene-editing enzyme inadvertently modifies genomic DNA sequences other than the intended target. This could potentially lead to mutations that disrupt normal gene function, potentially causing harmful consequences such as the activation of oncogenes or the inactivation of tumor suppressor genes, which could increase the risk of cancer. One approach to this problem lies in improving the precision of gene editing tools. As the most-commonly used gene editing class, substantial effort has gone into improving the specificity of CRISPR-family proteins.^4–6^ Similar efforts are underway for meganucleases, zinc finger nucleases, or TAL effector nucleases as well.^13–16^ Nevertheless, it is likely that even newer, high-fidelity enzymes will retain some degree of off-target activity, and thus additional complementary approaches are needed to minimize the risks of their *in vivo* use, and to ensure the safety and efficacy of therapeutic applications.

An attractive and complementary strategy is to limit expression of gene editing enzymes (to the greatest degree possible) to the particular tissue of interest.^4–6^ Restricting the activity of gene editing enzymes to specific tissues or cell types can minimize off-target effects and reduce the risk of unintended genetic modifications in non-target tissues, thereby leaving healthy cells unaffected and improving the overall safety profile of gene therapies. Some degree of tissue specificity can be achieved through the choice of an appropriate vector. For example, certain AAV serotypes have preferential tropism for specific tissues,^17^ and substantial work has gone into developing lipid nanoparticles favoring delivery to particular body sites.^18^ Unfortunately, the tissue selectivity that has so far been achieved by either AAV or lipid nanoparticles is not absolute.^17,18^ This can be especially problematic for specific tissues known to be difficult to transduce *in vivo*, which often require the use of vectors with broad tropism. In such circumstances, an alternative approach is to limit expression of the transgene encoding the gene editing enzyme through genetic means, for example by using tissue-selective promoters or microRNA binding sites.^19,20^

Here we evaluate the use of these approaches in the context of a gene therapy targeting latent HSV harbored within sensory ganglionic neurons, with the therapeutic gene editing enzyme being delivered by the commonly used and broadly tropic AAV serotype AAV9.^21^ Our results demonstrate that the inclusion of either tissue-selective promoters or microRNA binding sites can markedly reduce the occurrence of off-target effects, while retaining good gene editing efficacy at the target tissue. These results argue for consideration of tissue-selective transgene expression during the development and implementation of *in vivo* gene editing therapeutics.

## RESULTS

### GUIDE-tag identifies potential off-targets of Cas9/sgPCSK9 in NIH3T3 cells

In our previous studies, we utilized adeno-associated virus (AAV)-mediated delivery of the m4 meganuclease to efficiently eliminate latent HSV-1 genomes from ganglionic neurons in mice. The recombinant m4 meganuclease specifically recognizes and cleaves a target sequence located within *ICP0*, a duplicated gene in HSV-1 genome. More than 90% of latent HSV genomes in the ganglia of mice was eliminated using AAV-mediated delivery of m4 transgene under a ubiquitous CBh promoter (AAV9-CBh-m4).^22^ However, unintended adverse effects were also observed in those mice. Mice administered AAV9-CBh-m4 exhibited clinical signs of hepatotoxicity, including weight loss, bloating, and general health decline.^21^ These adverse clinical effects may be due to overexpression of the m4 transgene in liver, as has been observed with a variety of other gene therapies utilizing AAV delivery.^23^ In addition to these acute effects, the overexpression of m4 in liver raises the possibility of off-target gene editing in this non-relevant tissue, which is outside the scope of our therapeutic target of HSV.

To identify potential m4 off-target sites, we first employed GUIDE-tag analysis,^24^ which is analogous to GUIDE-seq, a gold standard for detecting off-target cleavage sites.^25^ Both techniques rely on the integration of double-stranded oligodeoxynucleotides (dsODNs) at double-strand breaks (DSBs) induced by DNA nucleases. However, unlike GUIDE-seq, which requires DNA shearing during next-generation sequencing (NGS) library preparation, GUIDE-tag utilizes tagmentation. Tagmentation combines DNA fragmentation and adapter ligation steps into a single, efficient process, reducing time and complexity. To validate the GUIDE-tag method, we conducted experiments using NIH3T3 mouse fibroblast cells, targeting a well-characterized promiscuous site in the *PCSK9* gene. This site has been frequently used in prior studies for the comparison of off-target detection strategies.^24,26,27^ NIH3T3 cells were co-transfected with pX330-PCSK9, which encodes SpCas9 and a single-guide RNA (sgRNA) targeting *PCSK9*, along with blunt iGUIDE donor, the longer dsODN designed to minimize mis- priming artifacts.^28^ The 5ʹ ends of the dsODN were phosphorylated, and phosphorothioate linkages were incorporated at both the 5ʹ and 3ʹ ends of each strand to enhance the stability of the dsODN in cells. Following transfection, GUIDE-tag analysis was performed, and an NGS library was prepared for sequencing. The NGS sequencing yielded approximately 1.5 million sequencing reads, from which 228,583 unique molecular identifiers (UMIs) were identified. We identified 68 off-target sites where the UMI counts exceeded 0.2% of the total UMI population (Figure 1A). Most off-target sites contained one or two mismatches within the protospacer and/or utilized an alternative PAM sequence, such as NAG. Among the off-target sites of Cas9/PCSK9 identified in NIH3T3 cells, 23 overlapped with the 40 off-target sites reported in the GUIDE-tag analysis of mouse liver,^24^ and 20 matched the 26 off-target sites identified by the DISCOVER-seq analysis of murine B16-F10 cells.^27^ Additionally, we identified 44 novel off-target sites that were not detected by earlier methods (Figure 1B). The observed differences in off-target site detection by GUIDE-tag for Cas9/PCSK9 between mouse liver^24^ and our study are likely due to epigenomic differences in the host cell genomes or other technical factors. This result indicates that GUIDE-tag analysis effectively detects both on-target and off-target sites of CRISPR/Cas9 in NIH3T3 cells, and thus can be applied to identify potential m4 off-target sites.

**Figure 1.**
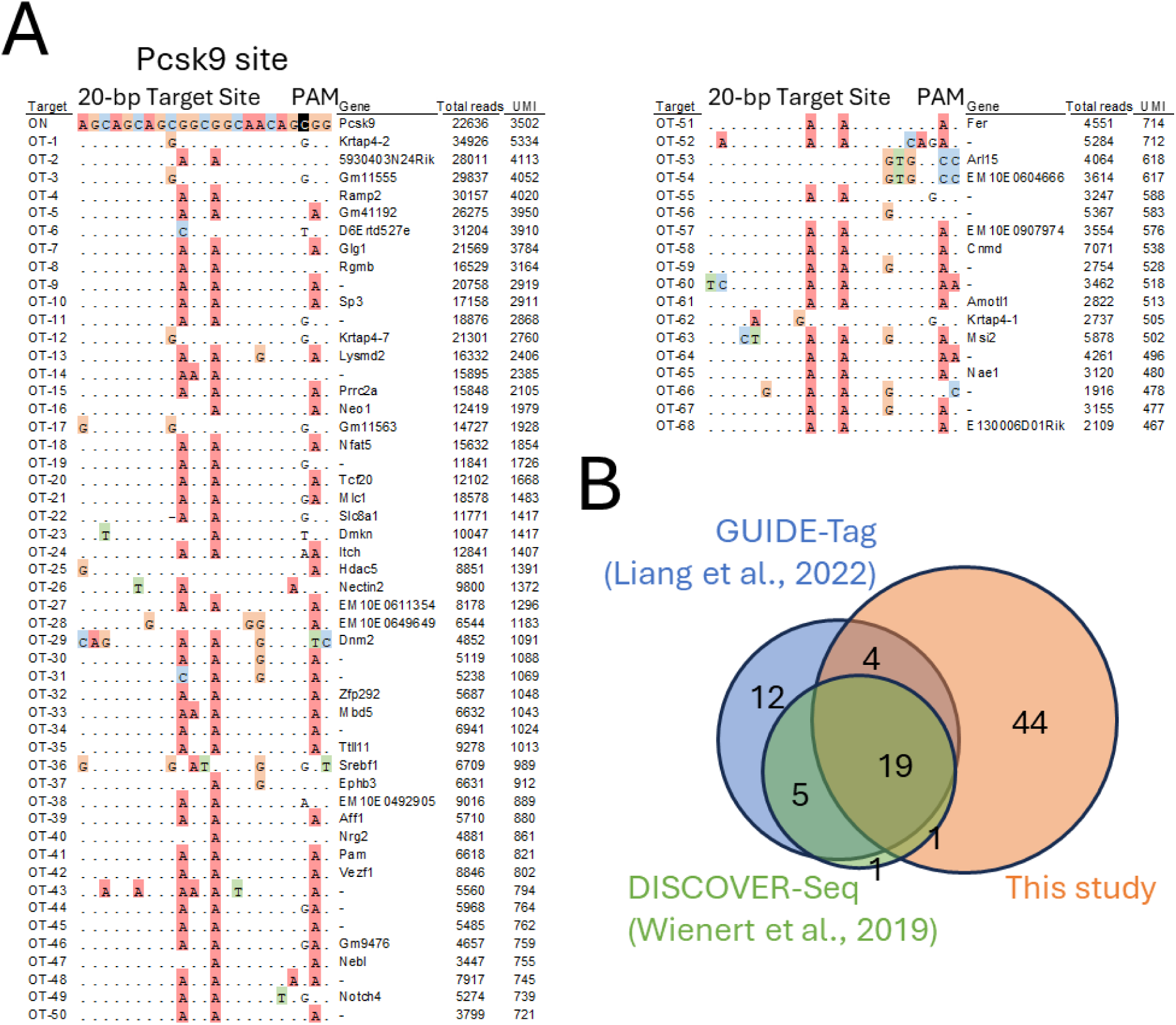
Off-target sites of the Cas9/sgPCSK9 in NIH3T3 cells. (A) Off-target (OT) sites of Cas9/sgPCSK9 in NIH3T3 cells identified by GUIDE-Tag. NIH3T3 cells were transfected with plasmid expressing Cas9 and a promiscuous sgRNA targeting PCSK9 (pX330-PCSK9) along with an iGUIDE donor. Off-target sites were discovered using GUIDE-Tag analysis of isolated DNA. Off-target sites with UMI counts exceeding 0.2% of the total UMI reads are listed. Mismatches relative to the on-target site are highlighted in color, and the gene name, total reads, and UMI counts for each site are shown. (B) Venn diagram of the overlap of off-target sites for sgPCSK9 identified in previous studies using GUIDE-Tag (40 loci) and Discover-Seq (26 loci), in comparison with the 68 loci identified in the current study.

### GUIDE-tag identifies potential off-target sites for m4

For the GUIDE-tag analysis of meganuclease m4, we first established NIH3T3 cells containing the on-target sequence of m4 (NIH3T3-m4TS) within their genome through lentiviral transduction followed by puromycin selection. NIH3T3-m4TS cells were then co-transfected with the plasmid DNA encoding m4 (CMV-m4) and iGUIDE donor with 4-nucleotide (nt) 3’ overhangs on both ends. The iGUIDE donor was designed to specifically integrate at the m4 cleavage site and its off-target sites. As an engineered variant of the I-CreI meganuclease,^29,30^ m4 cleaves double-stranded DNA, generating 4-nt overhangs with 3’-OH termini.

Integration sites of the dsODN within the NIH3T3-m4TS genome were mapped using NGS. From more than 1.8 million sequencing reads, 230,906 UMIs were identified. The analysis revealed 54 off-target sites where UMI counts exceeded 0.2% of the total UMI population (Figure 2A). We then compared the UMI counts of the top 100 off-targets identified by GUIDE-tag analysis between Cas9/PCSK9 and m4, relative to the on-target UMI counts. Interestingly, the relative UMI counts of m4 off-targets are lower than that of Cas9/PCSK9, suggesting overall better specificity (Figure 2B). However, compared to the sequence mismatch level of Cas9/PCSK9 off-targets, which averaged 3.4 mismatches within a 23-base pair target sequence, m4 off-targets displayed a significantly higher mismatch level, averaging 9.5 mismatches within a 24-base pair target sequence (Figure 2A). These results indicate that m4 is capable of cleaving DNA sequences containing more than 9 mismatches to the target sequence, though at a relatively low probability. Unlike CRISPR-Cas9, which achieves DNA recognition specificity through the formation of an RNA-DNA heteroduplex, the protein-DNA binding affinity and specificity of meganuclease are influenced by multiple factors, including the protein structure, the interface of protein-DNA interactions, and the extent of DNA bending by protein binding.^13^

**Figure 2.**
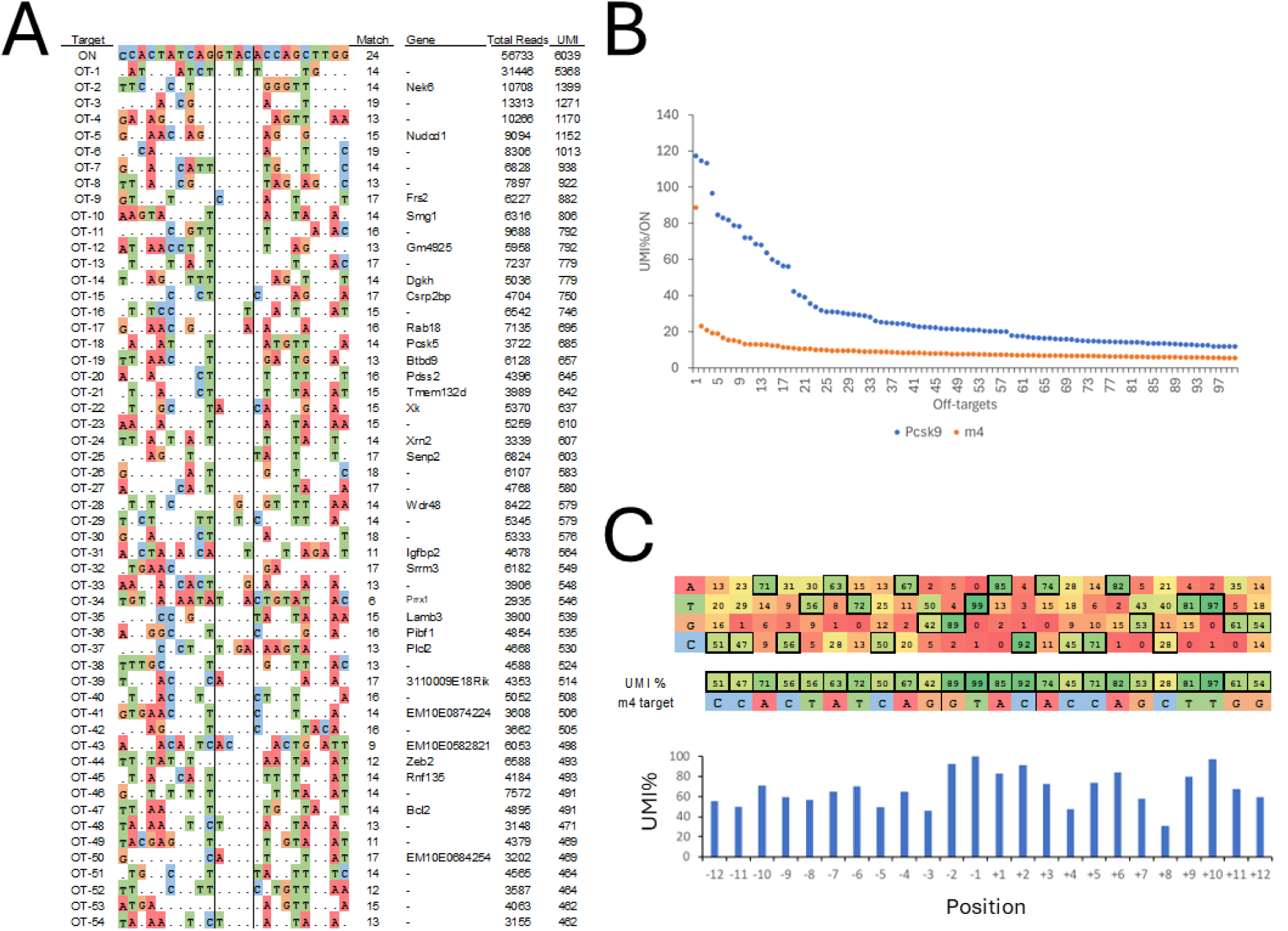
Off-target sites of the meganuclease m4 in NIH3T3-m4TS cells. (A) Off-target sites identified by GUIDE-Tag in NIH3T3-m4TS cells. NIH3T3-m4TS cells (NIH3T3 cells with the m4 target sequence integrated using lentiviral transduction) were transfected with a plasmid expressing m4 (CMV-m4) and an iGUIDE donor with 3’-overhang dsDNA sequences in both ends, designed to integrate into genomic DNA cleaved by m4. Off- target sites with UMI counts greater than 0.2% of the total UMI reads are listed. Mismatches compared to the on-target site of m4 are highlighted in color, along with the gene name, total reads, and UMI counts for each site. The central four nucleotides are positioned between horizontal lines. (B) Comparison of relative off-target UMI counts for GUIDE-Tag between Cas9/sgPCSK9 and m4. The UMI counts of the top 100 off-target sites relative to the on-target UMI counts are shown. (C) Identification of the cleavage specificity of m4 at individual positions within the DNA target site by GUIDE-Tag analysis. The nucleotides of m4 on-target sequences are boxed. The percentages of UMIs at individual positions for the on-target site and 54 off- target sites of m4 were analyzed.

To evaluate the sequence specificity of individual bases within the m4 DNA target sequence, UMI percentages at each nucleotide position were calculated using UMI counts from both the on-target site and the 54 identified off-target sites. m4 exhibited high specificity at the central four nucleotides, as well as at positions +6, +9, and +10, while exhibiting notable promiscuity at positions +8, -3, +4, and -11 (Figure 2C). These findings provide crucial information for enhancing the fidelity of m4 through the mutagenesis of amino acids that interact with these nucleotides.

### Expression of m4 from a ubiquitous promoter results in a high level of off-target gene editing in liver

The potential m4 off-target sites identified through GUIDE-tag analysis in NIH3T3 fibroblast cells require validation in an animal model to ensure the feasibility and safety of the therapeutic approach. In addition, since GUIDE-tag identifies DSBs rather than mutagenesis, and not all DSBs are converted to gene editing,^31^ *in vivo* assessment of genetic alterations at the identified off-target sites is necessary.

We evaluated the editing rates of m4 off-targets using DNA samples from the liver (n=2) and trigeminal ganglia (TG, n=4) of mice administered AAV9-CBh-m4 as well as control mice (n=2). TG is the primary target tissue for our HSV therapy, as it is the tissue where latent HSV resides after ocular infection. On the other hand, the liver, although readily transduced by AAV9, is not relevant to latent HSV disease. To minimize experimental complexity, these mice were not infected with HSV. In AAV9-CBh-m4 transduced cells, m4 expression is driven by the ubiquitous CBh promoter, a hybrid of the cytomegalovirus (CMV) enhancer and chicken β-actin (CBA) promoter.^32^ The editing rates of the top 20 off-targets identified by GUIDE-tag analysis were determined through amplicon deep sequencing (Figure S1), and 10 of these off-targets showed more than 5% editing in at least one mouse DNA sample (Figure 3A). AAV loads in the samples were quantified by ddPCR (Figure 3B).

**Figure 3.**
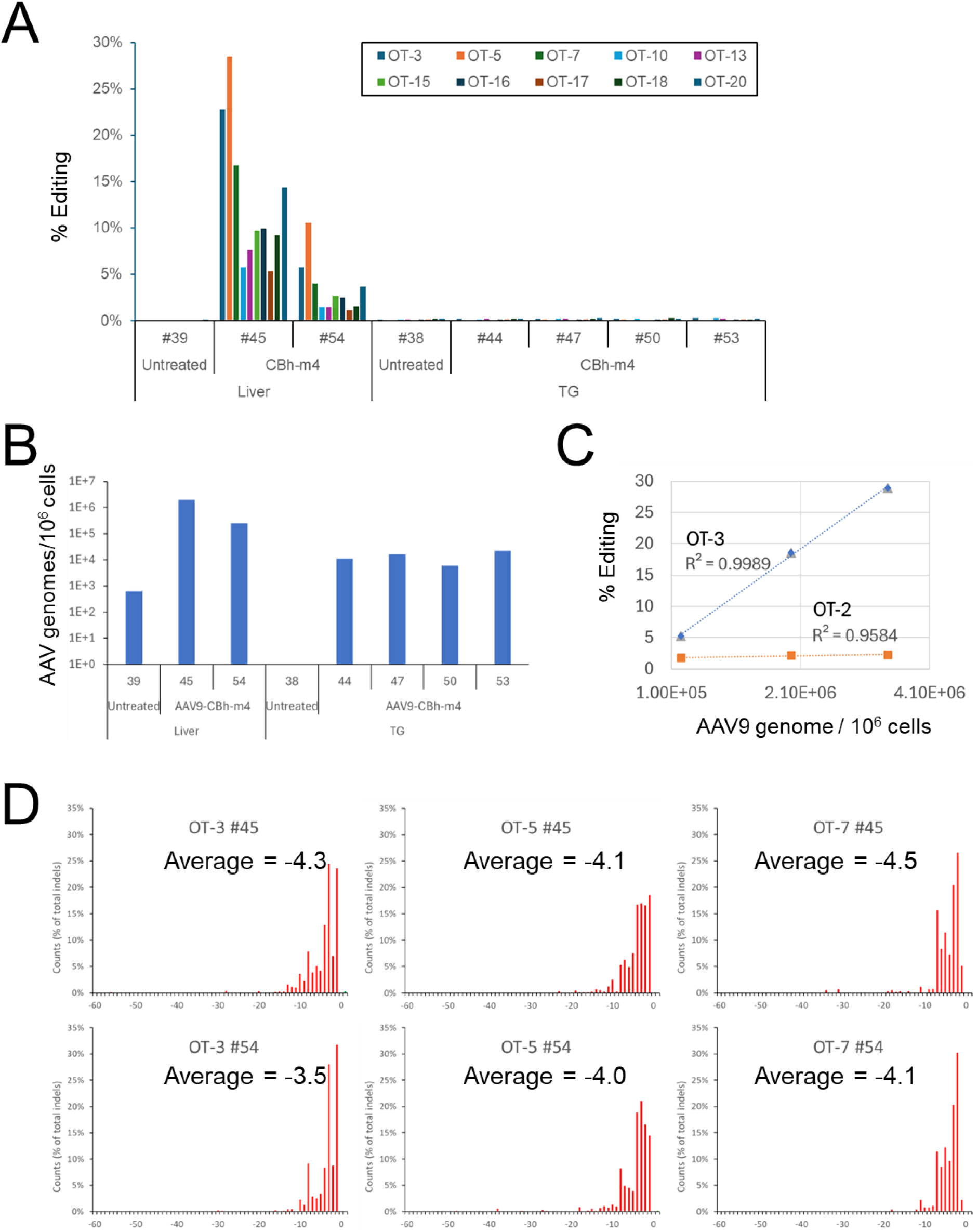
Validation of off-target sites for meganuclease m4 in liver and trigeminal ganglia (TG) tissue DNA from mice infected with AAV9-CBh-m4. (A) Detection of off-target editing rates in liver or TG tissue DNA using amplicon deep sequencing. Editing rates of the 10 off-target sites identified by GUIDE-Tag were determined through amplicon sequencing of liver (n=2) and TG (n=4) tissue DNA from mice infected with AAV9-CBh-m4, which expresses m4 driven by a ubiquitous CBh promoter. (B) AAV loads. AAV loads in the liver or TGs of mice infected with AAV9-CBh-m4 were quantified using ddPCR. (C). Correlation between AAV loads and editing rate. AAV loads and editing rates of OT-2 and OT-3 in liver tissue DNA from mice infected with AAV9-CBh-m4 (n=3) were analyzed. (D) The frequency distribution of indel sizes at off-target sites of m4. The relative frequency of indel sizes at three off-target sites (OT-3, OT-5 and OT-7) of m4 was determined by amplicon sequencing of two liver tissue DNA from mice infected with AAV9-CBh-m4 and CRISPResso2 analysis of base editing data.

The editing rates of 4 off-targets (OT-3, OT-5, OT-7, and OT-20) in liver DNA from AAV9-CBh- m4 treated mice were high, ranging from 14.33% to 28.49% in mouse #45 and 3.65% to 10.57% in mouse #54. Amplicon deep sequencing of OT-2 and OT-3 was repeated using three liver tissue DNA from AAV9-CBh-m4 treated mice. Interestingly, the editing rates were positively correlated with AAV loads, with correlation coefficients of R = 0.9584 and R = 0.9989 for OT-2 and OT-3, respectively (Figure 3C). The editing rates of these off-target sites in TG DNA from AAV9-CBh-m4 treated mice were substantially lower, with a maximum of 0.29% (Figure 3A). The high editing rates of the four off-targets in liver are likely due at least in part to the high level of AAV9 virus transduction in this tissue, since liver tissue showed approximately 2 logs higher AAV9 viral load than TG (Figure 3B). The deletion sizes, as detected by amplicon deep sequencing of off-target sites OT-3, OT-5, and OT-7, ranged from 1 to 10 base pairs, with an average size of 4 base pairs (Figure 3D). This result demonstrates that m4 expression, when driven by a ubiquitous promoter, results in a high level of off-target gene editing in liver. Because HSV-1 does not establish latency in liver, the expression of m4 in the liver would provide no clinical benefit, and thus we sought to evaluate approaches to minimize m4 expression in liver tissues.

### Use of a neuron-specific promoter reduces off-target effects in liver

In our previous studies, we restricted m4 expression to neuronal tissues to minimize the liver-dependent adverse clinical effects observed in mice treated with AAV9-CBh-m4.^21^ We used AAV9-E/CamKII-m4, which utilizes the neuron-specific E/CamKII, which is comprised of the Calmodulin Kinase II (CamKII) promoter^33^ combined with the CMV enhancer (E). Unlike mice administered AAV9-CBh-m4, mice treated with AAV9-E/CamKII-m4 showed minimal or no clinical symptoms or pathological findings in the liver.^21^

The difference in clinical outcomes observed in the livers of AAV9-CBh-m4- and AAV9- E/CamKII-m4-treated mice suggested that neuron-specific expression might also reduce off-target gene editing within the liver. To examine these differences, we compared the genetic alterations in liver tissue samples from AAV9-E/CamKII-m4-treated mice (n=2) to those from AAV9-CBh-m4-treated mice (n=2). (Figure 4A). We performed amplicon deep sequencing to assess the genetic alterations of the 10 m4 off-targets mentioned above. Though AAV loads in the liver were similar (Figure 4B), AAV9-E/CamKII-m4-treated mice demonstrated a significantly reduced level of genetic alterations in the liver compared to AAV9-CBh-m4-treated mice. (Figure 4A). For instance, liver DNA from AAV9-E/CamKII-m4-treated mice showed editing rates for four off-targets (OT-3, OT-5, OT-7, and OT-20) ranging from 0.03% to 0.41%, significantly lower than the 4.01% to 28.49% found in AAV9-CBh-m4-treated mice (Figure 4A). Indel formation caused by m4 expression was detected through multiple sequence alignments (Figure 4C). These results suggest that restricting meganuclease expression to neuronal tissues via the E/CamKII promoter reduces off-target effects in the liver, thereby improving the safety of meganuclease expression *in vivo*.

**Figure 4.**
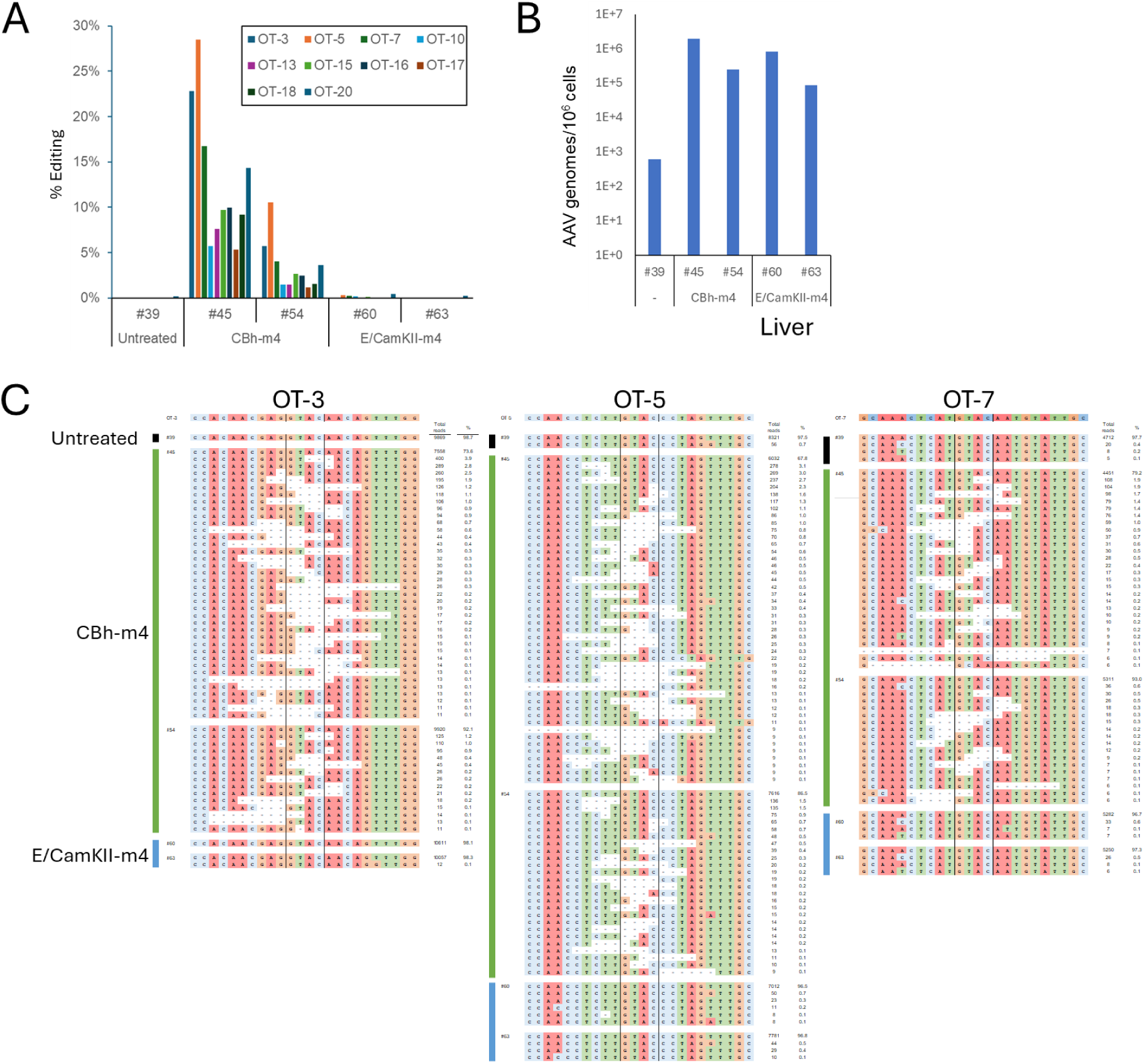
Reduction of off-target editing in the liver by expression of m4 from the neuron- specific promoter CamKII. (A) Detection of off-target editing in liver tissue DNA from mice infected with AAV9-CBh-m4 or AAV9-E/CamKII-m4 using amplicon deep sequencing. Editing frequencies of the 10 off-target sites identified by GUIDE-Tag were determined through amplicon sequencing of liver tissue DNA from mice infected with AAV9-CBh-m4 (n=2) or AAV9-E/CamKII-m4 (n=2). (B) Comparison of AAV loads. AAV loads in the liver of mice infected with AAV9-CBh-m4 or AAV9-E/CamKII-m4 were quantified using ddPCR. AAV9-E/CamKII-m4 expresses m4 driven by the neuron-specific CamKII promoter with portions of CMV enhancer. (C) Decreased off-target indel formation in the liver by the expression of m4 driven by neuron-specific promoter. Indel sequences and their frequencies at the OT-3, OT-5, and OT-7 sites were determined by amplicon sequencing of liver tissue DNA from mice infected with AAV9-CBh-m4 or AAV9-E/CamKII-m4 and CRISPResso2 analysis of base editing data.

### Meganuclease expression driven by the E/CamKII promoter also induces fewer off-target effects in TG compared to the CBh promoter

We also compared the genetic alterations in the TG, the relevant target tissue for anti-HSV therapy, between AAV9-E/CamKII-m4- and AAV9-CBh-m4-treated mice (Figure 5). In our previous study, the AAV9-E/CamKII-m4 regimen demonstrated significantly improved tolerability, including decreased hepatotoxicity and neurotoxicity, compared to AAV9-CBh-m4.^21^ However, the reduction in HSV-1 viral load was slightly lower in AAV9-E/CamKII-m4-treated mice compared to AAV9-CBh-m4-treated mice in both the superior cervical ganglia (SCG) and TG.^21^ We performed amplicon deep sequencing of the 10 m4 off-targets using TG DNA samples from AAV9-E/CamKII-m4-treated mice (n=4) and compared the results to those from AAV9-CBh-m4- treated mice (n=4) (Figure 5A). AAV loads in the TG of these mice were comparable (Figure 5B). Intriguingly, AAV9-E/CamKII-m4-treated mice also appeared to exhibit slightly lower levels of off-target editing rates in the TG compared to AAV9-CBh-m4-treated mice, with levels approaching those of the control mice. For example, TG DNA from AAV9-E/CamKII-m4-treated mice showed an average editing rate of 0.19% for OT-20, comparable to control, whereas the editing rate in AAV9-CBh-m4-treated mice was 0.24%.

**Figure 5.**
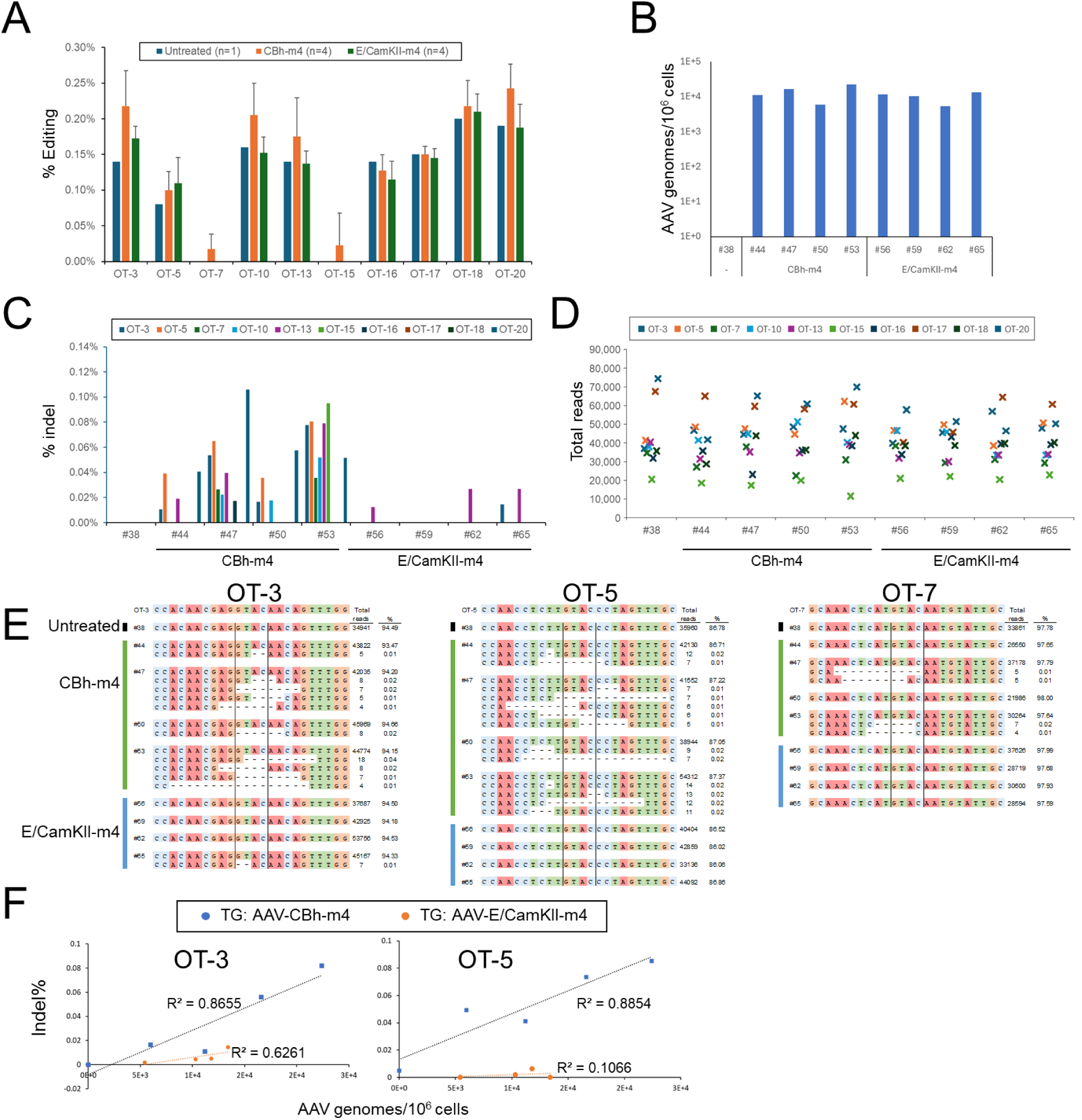
Comparison of off-target editing in the TG DNA by m4 driven by CBh or E/CamKII promoters. (A) Detection of off-target editing in TG tissue DNA using amplicon deep sequencing. Editing rates of the 10 m4 off-target sites identified by GUIDE-Tag were determined through amplicon sequencing of TG tissue DNA from mice infected with AAV9-CBh-m4 (n=4) or AAV9-E/CamKII- m4 (n=4). (B) Comparison of AAV loads in TG. AAV loads in the TG of mice infected with AAV9- CBh-m4 or AAV9-E/CamKII-m4 were quantified using ddPCR. (C) Decreased off-target indel formation in TG DNA by the expression of m4 driven by neuron-specific promoter. Indel frequencies at the OT-3, OT-5, and OT-7 sites were analyzed through amplicon sequencing of TG tissue DNA from mice infected with AAV9-CBh-m4 or AAV9-E/CamKII-m4. (D) The total read numbers from amplicon sequencing of off-target sites are presented. (E) Indel formation in the TG tissue DNA by the expression of m4 driven by CBh promoter or E/CamKII promoter. Indel percentile at OT-3, OT-5, and OT-7 were analyzed through amplicon sequencing of TG tissue DNA from mice infected with AAV9-CBh-m4 or AAV9-E/CamKII-m4, followed by CRISPResso2 analysis of base editing data. (F) Correlation between AAV loads and indel formation. AAV loads and indel percentile of OT-3 and OT-5 in TG tissue DNA from mice infected with AAV9-CBh-m4 or AAV9-E/CamKII-m4 were analyzed.

The editing rates observed in our assay include the deep sequencing error rate. The error rate of conventional NGS ranges between 0.1% to 1%.^34^ Since the majority of NGS errors are nucleotide substitutions, we chose to also analyze the indel percentage in amplicon deep sequencing results (Figure 5C). Although the total read numbers between these samples were comparable (Figure 5D), analysis of TG DNA from AAV9-CBh-m4-treated mice showed up to 0.106% of indel formation. In contrast, TG DNA from AAV9-E/CamKII-m4-treated mice identified indel percentage of up to 0.027%, lower than that of AAV9-CBh-m4 (Figure 5C and 5E). The editing rates of OT-3 and OT-4 in TGs positively correlated with AAV loads in AAV9-CBh-m4- treated mice, with correlation coefficients of R = 0.8655 and R = 0.8854, respectively (Figure 5F). However, no significant correlation was observed between editing rates and AAV loads in TG tissues from AAV9-E/CamKII-m4-treated mice.

### Restriction of meganuclease expression through a combination of neuron-specific promoter and liver-specific miRNA target sequence leads to additional reduction of off-target editing in liver and TG

Restricting meganuclease m4 expression using the neuron-specific E/CamKII promoter significantly reduced off-target effects in the liver. However, the genetic alteration levels of off-target sites (ranging from 0.08% to 0.57% in OT-3, OT-5, and OT-7) were still at a level that could be concerning for clinical application (Figure 4B). The genetic alterations observed in the liver of AAV9-E/CamKII-m4-treated mice might be due to the leaky expression restriction of E/CamKII.

In addition to using tissue-specific promoters to confine transgene expression to target tissues, the incorporation of microRNA (miRNA) target sequences has been shown to suppress transgene expression in non-relevant target tissues.^35^ miRNAs bind to specific target sequences in the 3’ untranslated regions (3’UTR) of mRNAs, inducing either mRNA degradation or translational repression.^36^ We first examined whether liver-specific miRNA-mediated repression of m4 expression could decrease liver toxicity (Figure 6A). We created AAV9-CBh-m4 with or without a 3X repeat of the miRNA-122 target sequence in the 3’UTR.^37^ miRNA-122 is known to be a regulator of fatty-acid metabolism and predominantly expressed in the liver,^38^ with minimal or undetectable levels in neuronal cells under normal physiological conditions. We observed that latently infected mice treated with a high dose (2 × 10^12^ vg) of AAV9-CBh-m4-miR122TS showed similar weight change with AAV9-CBh-m4 treated mice (Figure 6B), along with a reduction of ganglionic viral loads in SCG and TG of HSV1 infected mice (Figures 6C and 6D). AAV loads in the SCG, TG, and liver were comparable between the AAV9-CBh-m4 and AAV9- CBh-m4-miR122TS treated mice (Figures 6E, 6F, and 6G). Notably, mice treated with AAV9- CBh-m4-miR122TS displayed reduced histopathological signs of toxicity in the liver (e.g., formation of inflammatory cell foci), as well as in the TG (e.g., inflammation and axonopathy) compared to those treated with AAV9-CBh-m4 (Figures 6H, 6I, and 6J). This indicated that the AAV9-CBh-m4-miR122TS regimen maintains its anti-HSV efficacy while demonstrating improved tolerability compared to AAV9-CBh-m4.

**Figure 6.**
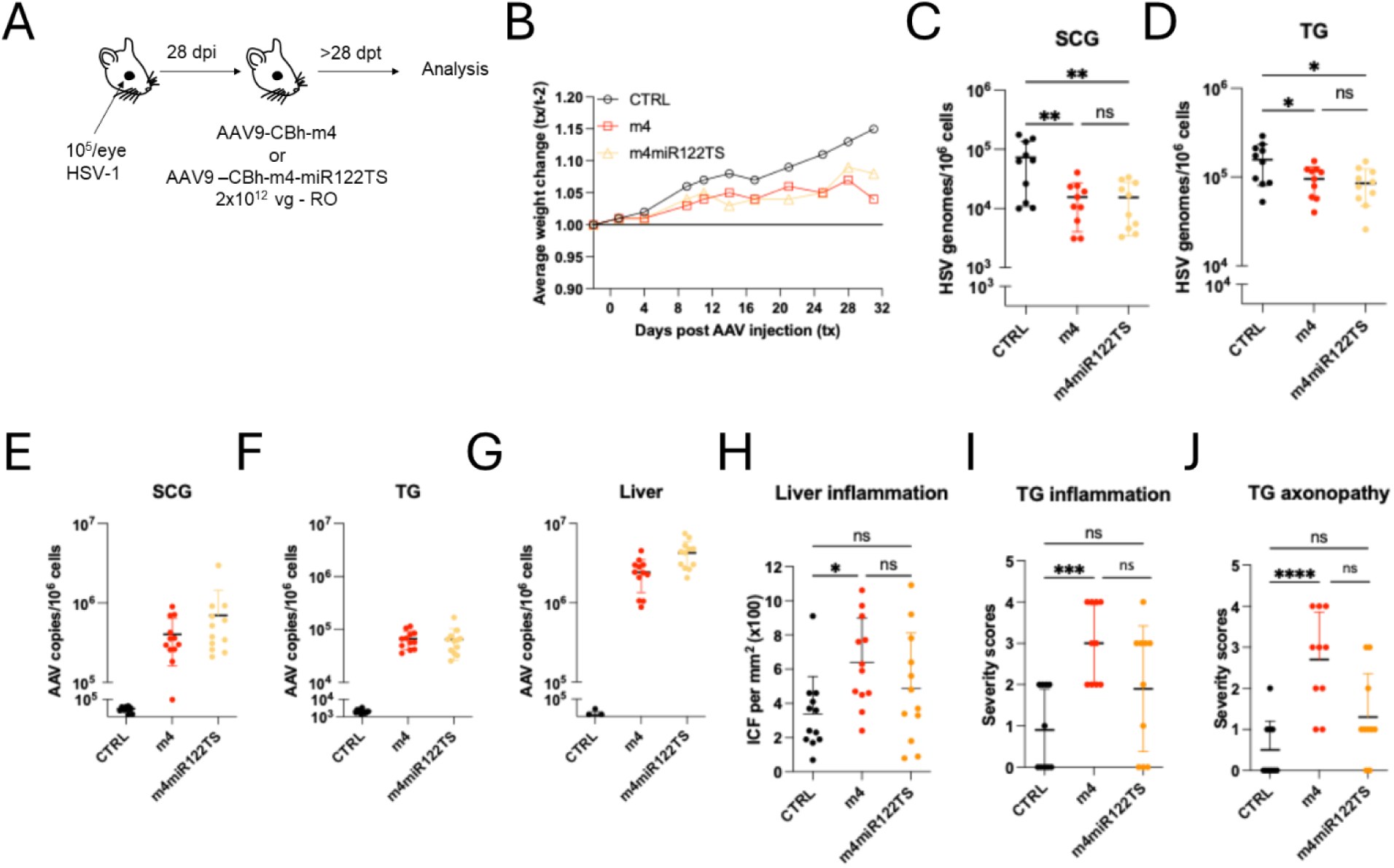
Efficacy and toxicity of AAV9-CBh-m4 with miRNA-mediated inhibition of liver transgene expression. (A) Experimental timeline of ocular infection, and meganuclease treatment administered by retro-orbital (RO) injection. (B) Average weight change after AAV administration in control infected mice (CTRL, n = 10), and infected mice treated with meganuclease m4 either without miR122TS (m4, n = 10) or with miR122TS (m4-miR122TS, n = 10), (C-D) Percent decrease of HSV loads in treated mice compared to control mice in SCG (C) and TG (D) and statistical analysis (ordinary one-way Anova, multiple comparisons with ns: not significant, *: *p* < 0.05, **: *p* < 0.01) are indicated. (E-G) AAV loads in SCGs (E), TGs (F) and liver (G) from control infected mice and infected mice treated. (H) Inflammatory cell foci (ICF) in liver sections from either HSV infected control mice or mice treated with m4 or m4-miR122TS (p < 0.05 for m4) with statistical analysis (Ordinary one-way Anova, multiple comparisons with ns: not significant; *: *p* < 0.05; \*\*\**p* <0.001; ****: *p* < 0.0001). (I-J) Severity scores of inflammation (H; p = <0.0001 for m4) and axonopathy (I; p = <0.0001 for m4) in TG from infected control mice and infected mice treated with m4 or m4-miR122TS.

Next, we sought to determine whether the combination of a neuron-specific promoter and a liver-specific miRNA target sequence in the m4 expression vector could minimize off-target effects in non-relevant target tissues. We created AAV9-CBh-m4 and AAV9-E/CamKII-m4 vectors, with or without the 3x repeats of miRNA-122 target sequence in the 3’UTR (Figure 7A).

**Figure 7.**
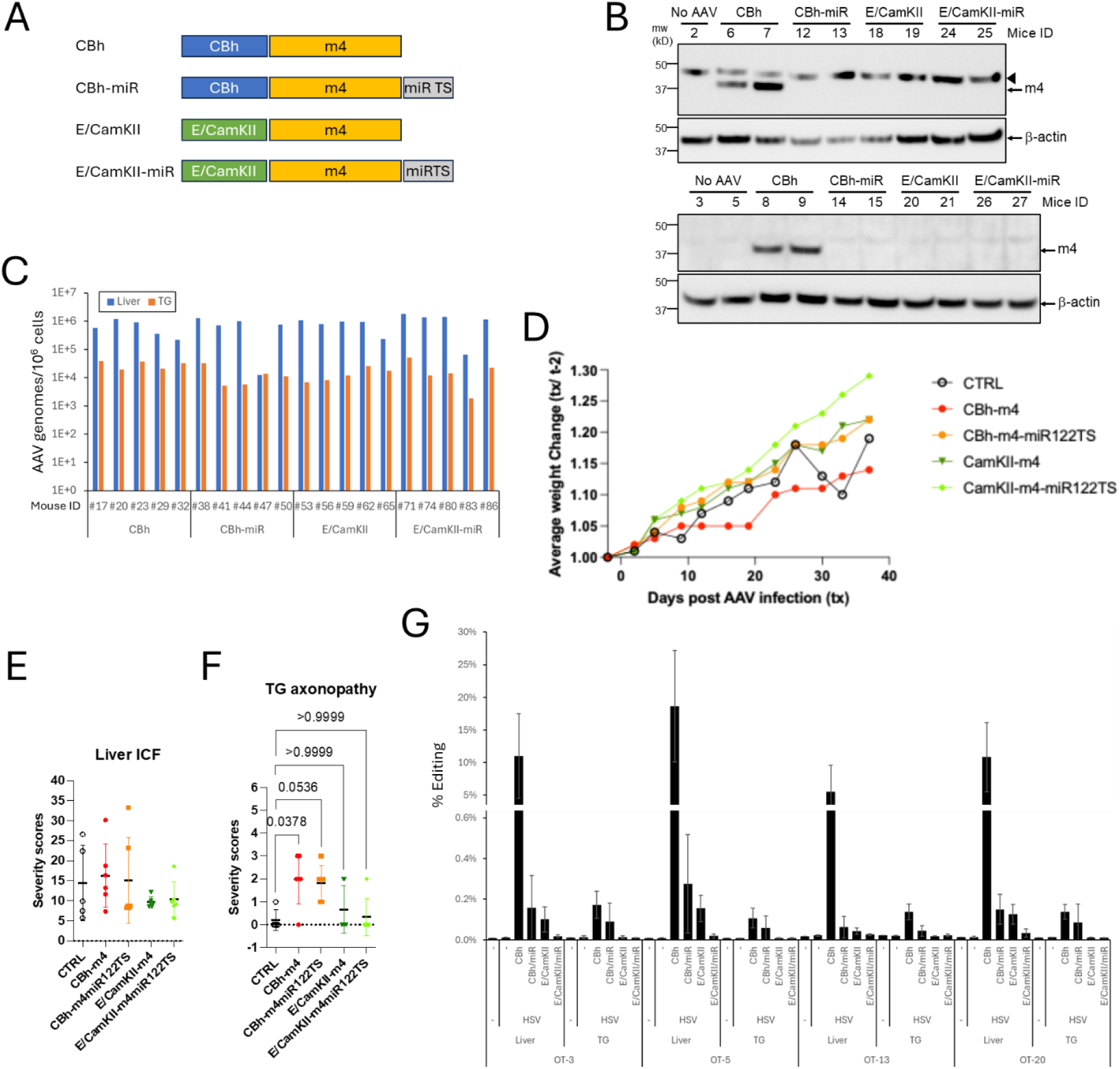
Combined reduction of off-target editing in liver and TG tissue DNA through regulated m4 on-tissue expression using a neuron-specific promoter and liver-specific miRNA. (A) Schematic representation of AAV constructs. (B) Western blot analysis of m4 expression (anti-HA) in livers from mice infected with AAV9-CBh-m4, AAV9-CBh-m4-miR122TS, AAV9- E/CamKII-m4, or AAV9-E/CamKII-m4-miR122TS, as well as control mice. Lysates were prepared and subjected to Western blot with anti-HA and anti-β-actin antibodies. The arrowhead indicates nonspecific bands. (C) Average weight change after AAV administration in control infected mice (CTRL, n = 10), and infected mice treated with meganuclease m4 either without miR122TS (m4, n = 10) or with miR122TS (m4-miR122TS, n = 10). (D) Comparison of AAV loads. AAV loads in the liver and TG tissue DNA from mice infected with AAV9-CBh-m4, AAV9-CBh-m4-miR122TS, AAV9-E/CamKII-m4, or AAV9-E/CamKII-m4-miR122TS were quantified using ddPCR. (E) Inflammatory cell foci (ICF) in liver sections from either HSV infected control mice or mice treated with m4. (F) Severity scores of axonopathy in TG from infected control mice and infected mice treated with m4 with statistical analysis (Ordinary one- way Anova, multiple comparisons with ns: not significant). (G) Significant reduction of off-target editing by m4 in the TG and liver DNA using a neuron-specific promoter and/or liver-specific miRNA target sequence in the AAV vector. Editing rates of OT-3, OT-5, OT-13 and OT-20 were determined through amplicon sequencing of liver and TG tissue DNAs from mice infected with AAV9-CBh-m4 (n=5), AAV9-CBh-m4-miR122TS (n=5), AAV9-E/CamKII-m4 (n=5), or AAV9-E/CamKII-m4-miR122TS (n=5) and control mice (n=1).

Western blot analysis of m4 protein expression in livers from mice infected with AAV9-CBh-m4, AAV9-CBh-m4-miR122TS, AAV9-E/CamKII-m4, or AAV9-E/CamKII-m4-miR122TS demonstrated that detectable m4 expression was observed only in livers from mice infected with AAV9-CBh-m4 (Figure 7B). This suggests that the inclusion of a liver-specific miRNA target sequence and/or a neuron-specific promoter effectively prevents m4 expression in non-relevant tissues with high AAV loads, particularly the liver. As expected, the liver tissue exhibited an AAV9 viral load approximately 2 logs higher than that in the TG (Figure 7C). Restricting m4 expression with either the E/CamKII promoter or miRNA122TS resulted in improved weight gain in treated animals, with the most robust weight gain being in receiving AAV9-E/CamKII-m4- miR122TS (Figure 7D). We found that restricting meganuclease m4 expression using the E/CamKII promoter was more effective in reducing histopathological signs of liver toxicity (e.g., inflammatory cell foci) and neurotoxicity in the TG (e.g., inflammation and axonopathy) than using miRNA122TS (Figures 7E and F).

We carried out amplicon deep sequencing of OT-3, OT-5, OT-13, and OT-20 on liver and TG DNA samples from mice treated with either AAV9-CBh-m4 (n=5), AAV9-CBh-m4-miR122TS (n=5), AAV9-E/CamKII-m4 (n=5), or AAV9-E/CamKII-m4-miR122TS (n=5), and control mice (n=1) (Figure 7G). As expected, the editing rates in liver DNA from AAV9-CBh-m4-treated mice were high, ranging from 5.54% to 18.62%. However, these percentages were substantially reduced with the use of the neuron-specific E/CamKII promoter (0.06% to 0.28%) or liver-specific miRNA-122 target sequence (0.04% to 0.15%). Notably, the combination of the E/CamKII promoter and miR122TS further reduced the editing rates to 0.02% to 0.03%, comparable to the control group (0.01% to 0.02%).

In TG DNA, AAV9-CBh-m4-treated mice showed editing rates between 0.11% and 0.17%. While AAV9-CBh-m4-miR122TS-treated mice showed a substantial reduction of editing rates in TG (0.04% to 0.09%), the levels were slightly higher than in the control group. However, the E/CamKII promoter alone was sufficient to bring the editing levels to those detected in the control group (0.01% to 0.02%).

These data indicate that the combined use of a neuron-specific promoter and a liver-specific miRNA target sequence in the 3’UTR can minimize off-target effects in the liver, one of the most critical non-relevant target tissues for potential off-target gene editing.

## DISCUSSION

Gene editing, along with related technologies such as base editing, prime editing, and gene writing, offer unprecedented opportunities in treating human disease.^39^ Gene editing approaches have shown impressive preclinical promise for treatment of a wide variety of inherited disorders, as well as for acquired conditions including cancer, transplant rejection, and infectious disease.^5^ Human clinical trials have been initiated for hereditary transthyretin amyloidosis (hATTR),^40^ hereditary angioedema (HAE),^41^ familial hypercholesterolemia,^42^ and HIV,^43^ among others. In addition, at least one gene editing therapy, Casgevy, for the treatment of Sickle cell disease and beta thalassemia, has been approved for clinical use in the UK and US,^8^

Some embodiments of gene editing involve *ex vivo* manipulation of an isolated cell type, such as engineering of CAR T cells for therapy of cancer or other disorders.^5^ For many applications, however, the most practical and/or efficacious use of gene editing would occur *in vivo*, meaning within the body of the affected individual.^4–6^ Like any *in vivo* gene therapy approach, effective gene editing requires efficient expression of the gene editing nuclease in the target tissue. This is often achieved through the use of gene therapy vectors with broad tissue tropism, often coupled with strong constitutive promoters. While such an approach may be warranted for some types of gene therapy, for gene editing this also risks the unnecessary exposure of non-target tissues to the gene editing nuclease.

The potential of gene editing enzymes to mediate off-target effects, meaning the induction of mutagenesis at genomic targets other than the desired site, is a well-known concern with gene editing approaches. Therefore, much effort has been expended toward the development of more-specific nucleases.^5,6^ The resulting nucleases appear to have substantially improved specificity relative to early-generation enzymes, although it is unlikely that off-target activity can ever be completely abolished. In any event, it remains unclear what degree of risk any residual levels of off-target gene editing might pose to individuals receiving gene editing therapy. To date, the limited trials of *in vivo* gene editing suggest that it is well tolerated and at least initially safe.^40–42^ Thus, the potential for off-target gene editing on the targeted tissue must be weighed against the potential beneficial effects of the therapy for the person.

As a general rule, it is desirable to limit exposure to any gene therapy modality, to the maximum degree possible, to the desired tissue, meaning the tissue where the therapeutic benefit can be realized. This is especially true for gene editing approaches, where potential off-target effects raise the possibility of mutagenesis and/or carcinogenesis in other tissues, especially tissues with inherently robust proliferative potential. In principle, the tissue tropism of the vector used for gene delivery could allow selective targeting of the desired target tissue, with minimal exposure of other tissue types. As a practical matter, however, there are certain tissues that are easily targeted by multiple vectors. For example, the liver is readily targeted by multiple AAV and lipid nanoparticle vectors.^17,44,45^ On the other hand, other tissues can be inherently much more difficult to target. A relevant example are neurons of the sensory ganglia, which are the relevant target for gene therapies for chronic pain, latent alphaherpesvirus infections, and other indications.^46^ These neurons have been a challenging target for gene therapy delivery, due to a combination of the need for vectors to cross the blood/peripheral nervous system barrier, the inherent limited transducibility of these cells, or other factors.^46^ Even relatively successful vectors for these applications, such as AAV9, transduce the liver to a higher degree than they do the desired ganglionic neurons.^21^ Thus, vector tropism alone is unlikely to fully solve the issue of off-target expression of gene therapy transgenes, especially when the desired target is not the liver.

An alternative approach to increase the selectivity of *in vivo* gene editing is the use of tissue-specific promoters. Such promoters are typically identified via the strong and tissue-selective expression of the genes they regulate in their native physiological state. Tissue specific promoters have been described for neurons,^33^ liver,^47^ pancreas,^48^ heart,^49^ and other tissues.^19^ However, tissue-specific promoters can have certain limitations that should be recognized. First, such promoters are rarely actually “tissue specific”; there is typically at least some level of observable expression in other tissues, and such promoters might be better thought of as “tissue selective”. For gene editing applications, the relative levels of promoter activity should be evaluated in the desired tissue vs. the most relevant off-target tissues. For the studies describe here, we demonstrated that the E/CamKII promoter allowed strong transgene expression and good gene editing efficiency in ganglionic neurons, while minimizing transgene expression and off-target nuclease activity in liver, arguable the most important off-target tissue in this setting.

Additionally, it should be noted that many tissue-specific promoters have weaker promoter activity in their target tissues than do constitutive promoters. In our studies, we found that the E/CamKII promoter led to modestly weaker nuclease expression in ganglionic neurons, and similarly weaker gene editing, compared with the CBh promoter used at the same AAV dose. In our opinion the modest drop in efficacy (which can be compensated for through increased AAV dose) is a reasonable trade-off for the marked reduction in off-target gene editing we observed in liver. However, such trade-offs must be considered on a case-by-case basis for each specific gene therapy application.

Another approach to improving the tissue specificity of gene therapy is via the incorporation of miRNA target sites within the 3’ untranslated region (UTR) of the transgene. MicroRNAs (miRNAs) are small, non-coding RNAs that enable post-transcriptional regulation of gene expression by binding to the 3’ untranslated regions (3’ UTRs) of target mRNAs, leading to their degradation or translational repression.^35^ miRNAs vary in their tissue expression patterns,^20^ and thus incorporation of the appropriate miRNA binding sites can be a powerful tool for achieving improved targeting of gene therapy. In this study we utilized a binding site for miRNA-122,^37^ which is highly expressed in liver tissue, based on previous data showing that liver toxicity was dose limiting for our AAV-mediated delivery of meganucleases.^21^ As predicted, incorporation of the miRNA-122 binding site into our vector dramatically improved the overall tolerability of our therapy. It had the further salutary effect of markedly reducing off-target gene editing within the liver, which although likely not directly dose-limiting, remains a concern during *in vivo* gene editing.

While our experiments were not designed to determine whether use of tissue-specific promoters or miRNA binding sites is superior in a general sense to maximize the tissue specificity of gene therapy, our results that either approach can effectively limit undesired transgene expression in liver, resulting in improved tolerability of our therapy and a marked reduction in off-target gene editing in liver. It is worth noting that tissue-specific promoters may provide a broader protective effect, since many miRNAs are highly restricted in their tissue expression, and thus any tissues not expressing the miRNA would presumably not be protected from undesired transgene effects. Combining the two approaches appeared to further increase the relative specificity of transgene activity, although additional experimentation would be required to fully understand the degree to which the combination of approaches may reduce expression in the target ganglionic neurons, and if so whether the additional reduction of off-target tissue expression warrants such a trade-off.

Other strategies can also help minimize off-target genome editing. For example, gene editing can be conditionally controlled by regulating the protein levels of gene-editing enzymes. This can be achieved by fusing the enzymes with a destabilizing domain (DD), which makes the fusion protein unstable and susceptible to rapid degradation via the ubiquitin-proteasome pathway. The addition of a specific small-molecule ligand binds to the DD, stabilizing the DD-fused protein and restoring the cellular levels of the gene-editing enzymes. ^50–52^ Another promising approach is epigenetic editing, which uses deactivated nucleases fused with transcriptional or epigenetic effector domains to enable precise and reversible modifications of the epigenome and/or chromatin structure. These methods induce heritable epigenetic changes that regulate gene expression without causing DSBs in the genome.^53,54^

The impetus of our work was to improve the tolerability of our ganglionic neuron-targeting gene editing approach, and at the same time to minimize off-target gene editing within tissues irrelevant to our therapeutic goals. The incorporation of either a neuron-specific promoter or the miRNA-122 binding site into our vectors appeared to readily achieve these goals. Nevertheless, it bears repeating that while reduction of off-target gene editing is a worthwhile goal on general principles, it remains unclear what if any deleterious effect low-level off-target gene editing may have in actual practice. The US FDA has recognized this gap in our current knowledge, and in 2022 released a draft guidance for “Human Gene Therapy Products Incorporating Human Genome Editing”,^1^ which was followed by a formal guidance document in January 2024.^55^ The current consensus is that the real-world consequences, if any, of off-target gene editing remain undefined, and as such, a maximum level of off-target activity consistent with regulatory approval has not been set at this point. Instead sponsors developing gene editing products are advised to evaluate on- and off-target activity in both the preclinical and clinical phases of development. The use of multiple methods is advised, including both genome-wide approaches and methods with adequate sensitivity to detect low-frequency events. Presumably, the hope is that over time such data will allow informed establishment of acceptable levels of off-target activity. Nevertheless, during this period, investigators in the field should continue work toward gene editing enzymes with improved specificity, as well as improved methods for restricting transgene expression to the therapeutically relevant tissues.

## MATERIALS and METHODS

### Generation of Plasmids

The pX330-PCSK9 vector was constructed by introducing a ‘G’ base and the *PCSK9* gene target sequence (AGCAGCAGCGGCGGCAACAG) into the pX330 vector (Addgene 42230). The lentiviral vector pLVX-EF1a-mTagBFP2-HSV1m4TS-eGFP, designed for the generation of NIH3T3-m4TS cells, was constructed using Gibson assembly by combining four DNA fragments: 1) the coding sequence of mTagBFP2, 2) the T2A peptide and m4 target sequence, 3) the coding sequence of EGFP, 4) the EcoRI/BamHI-digested pLVX-EF1α-IRES-Puro (Takara) backbone. The pCMV-m4 vector was created by replacing the *vsv-g* coding sequence in the pMD2.G vector with a hemagglutinin (HA) epitope tagged and nuclear localization signal (NLS)- linked m4 coding sequence, utilizing In-Fusion Advantage PCR cloning kit (Takara).

### Cell Culture and Transfection

NIH3T3 cells were cultured in Dulbecco’s modified Eagle’s medium (DMEM, Thermo Fisher) supplemented with 10% BCS (Thermo Fisher). NIH3T3-m4TS cells were generated via lentiviral transduction using the pLVX-EF1α-mTagBFP2-HSV1m4TS-eGFP vector. For lentivirus production, 12 μg of pLVX-EF1α-mTagBFP2-HSV1m4TS-eGFP, 4.2 μg of pMD2.G (Addgene #12259), and 7.8 μg of psPAX2 (Addgene #12260) were transfected into 5 million Lenti-X 293T cells (Takara) using the TransIT-LT1 reagent (Mirus). Three days post-transfection, the culture medium was collected, filtered through a 0.45 μm syringe filter (Thermo Fisher), and concentrated using Amicon Ultra centrifugal filters (∼100K, Millipore). Lentiviral transduction was performed by incubating NIH3T3 cells with the prepared virus in the presence of 8 μg/mL polybrene. After three days, NIH3T3-m4TS cells were selected by treating the culture with 2 μg/mL puromycin.

Electroporation of NIH3T3 and NIH3T3-m4TS cells for GUIDE-Tag analysis was conducted using the Neon transfection system (Invitrogen) following the manufacturer’s protocol. For every 50,000 cells, 0.5 μg of plasmid DNA (pX330-PCSK9 or pCMV-m4) was co-transfected with 80 pM of annealed dsODN. Three days after transfection, cells were harvested, and genomic DNA was extracted using the QIAamp DNA Kit (QIAGEN).

The dsODN used for GUIDE-tag analysis of Cas9/PCSK9 was prepared by annealing two oligos (IDTDNA). iGUIDE-F: /5Phos/G*C*TCGCGTTTAATTGAGTTGTCATATGTTAATAACGGTATACGC*G*A and iGUIDE-R: /5Phos/T*C*GCGTATACCGTTATTAACATATGACAACTCAATTAAACGCGA*G*C.

The dsODN for m4 was prepared by annealing following two oligos. iGUIDE-GTAC-F: /5Phos/G*C*TCGCGTTTAATTGAGTTGTCATATGTTAATAACGGTATACGCGAGT*A*C and iGUIDE-GTAC-R: /5Phos/T*C*GCGTATACCGTTATTAACATATGACAACTCAATTAAACGCGAGCGT*A*C. /5Phos/ indicates 5′ phosphorylation, and * denotes the presence of a phosphorothioate bond.

### GUIDE-tag Analysis

GUIDE-tag analysis was conducted as previously described,^24^ with minor modifications. For tagmentation, transposomes were assembled by incubating 1 μL of 100 μM annealed oligonucleotides (Tn5-A-bottom with either i5-N501 or i5-N502) and 1 μL of unloaded Tagmentase (Diagenode) at 23°C for 55 minutes. A mixture containing 200 ng of genomic DNA, tagmentation buffer (Diagenode), and 2 μL of assembled transposome in a final volume of 20 μL was incubated at 55°C for 7 minutes. The tagmented DNA was purified using a Zymo column (Zymo Research) and eluted in 22 μL of TE buffer.

The first PCR was performed using Platinum SuperFi DNA Polymerase, supplemented with final concentration of 0.5 M tetramethylammonium chloride. Two separate PCR libraries were generated using the following primer sets, i5 primer with either iGUIDE_insert_F or i5 primer and iGUIDE_insert_R (Table S1). The PCR products were cleaned with 1.2x reaction volume of SPRIselect beads (Beckman Coulter) and eluted in 10 μL of elution buffer (10 mM Tris–HCl, 0.1 mM EDTA). The second PCR was performed using the i5 primer and i7 indexed primers (N701 to N704). PCR products were purified with 0.9x reaction volume of SPRIselect beads to retain fragments larger than 150 bp. The final libraries were quantified using Tapestation and Qubit (Agilent) and sequenced on a MiSeq instrument (Illumina).

### Amplicon deep sequencing

Genomic DNAs were extracted from liver or TG tissues of AAV9-CBh-m4, AAV9-CBh-m4-miR, AAV9-E/CamKII-m4 or AAV9-E/CamKII-m4-miR using the DNeasy Blood and Tissue Kits (Qiagen). 100 ng of genomic DNA was used for PCR amplification with Phusion Plus DNA polymerase (Thermo Fisher) and locus-specific primers listed in Table S1. PCR products were purified using a 1.0X reaction volume of AMPure XP beads and eluted in 20 μL of DW. A total of 12 μL of DNA underwent processing with the Illumina DNA Prep (S) kit, followed by a second purification with a 1.0X reaction volume of AMPure XP beads and elution in 20 μL of DW. DNA concentration was quantified using the Tapestation and Qubit systems. The resulting amplicons were pooled at equimolar ratios and sequenced on a NovaSeq 6000 instrument (Illumina). Data from the amplicon sequencing were analyzed via CRISPResso2 (http://crispresso2.pinellolab.org/submission).

### Mice

Six- to eight-week old female Swiss Webster mice were purchased from either Taconic or Charles River, and housed in accordance with the institutional and NIH guidelines on the care and use of animals in research.

#### Ocular HSV infection

Mice were anesthetized by intraperitoneal injection of ketamine (100 mg/kg) and xylazine (12 mg/kg). Mice were infected in both eyes by dispensing 10^5^ PFU of HSV1 syn17+ contained in 4 uL following corneal scarification using a 28-gauge needle.

#### AAV inoculation

Mice anesthetized with isoflurane were administered the indicated AAV vector dose by unilateral retro-orbital (RO; ocular HSV infected mice) injection. Tissues were collected at the indicated time.

#### Study approval

All animal procedures were approved by the Institutional Animal Care and Use Committee of the Fred Hutchinson Cancer Center. This study was carried out in strict accordance with the recommendations in the Guide for the Care and Use of Laboratory Animals of the National Institutes of Health (“The Guide”).

### AAV production and titering

AAV vector plasmids pscAAV-CBh-m4, pscAAV-CBh-m4-miR122TS, pscAAV-E/CamKII-m4, and pscAAV-E/CamKII-m4-miR122TS were used to generate the AAV stocks in this study. These vectors express m4 meganuclease driven by either the ubiquitous CBh or neuron-specific E/CamKII promoter. An HA tag and NLS sequence was introduced upstream of the m4 gene, while three miR122 target sequence repeats were cloned downstream of the m4 gene.^37^ All AAV stocks were generated from transfected HEK293 cells and culture media produced by the Viral Vector Core of the Wellstone Muscular Dystrophy Specialized Research Center (Seattle). AAV stocks were generated by PEG-precipitation of virus from cell lysates and culture media, followed by iodixanol gradient separation^56,57^ and concentration into PBS using an Amicon Ultra-15 column (EMD Millipore). AAV stocks were aliquoted and stored at -80°C. All AAV vector stocks were quantified by qPCR using primers/probe against the AAV ITR, with linearized plasmid DNA as a standard, according to the method of Aurnhammer et al.^58^ AAV stocks were treated with DNase I and Proteinase K prior to quantification.

### ddPCR Quantification of viral loads in tissues

Total genomic DNA was isolated from ganglionic tissues using the DNeasy Blood and tissues kit (Qiagen, Germantown, MD) per the manufacturer’s protocol. Viral genomes were quantified by ddPCR in tissue DNA samples using an AAV ITR primer/probe set for AAV, and a gB primer/probe set for HSV as described previously.^59^ Cell numbers in tissues were quantified by ddPCR using mouse-specific RPP30 primer/probe set: Forward 5′- GGCGTTCGCAGATTTGGA, Reverse 5’- TCCCAGGTGAGCAGCAGTCT, probe 5’-ACCTGAAGGCTCTGCGCGGACTC. In some control ganglia, sporadic samples showed positivity for AAV genomes, although the levels were typically >2-3 logs lower than in ganglia from treated mice having received AAV. We attribute this to low-level contamination of occasional tissue samples.

### Western blot detection

Tissues lysates were obtained from approximately 20 mg liver tissue collected in 200 uL RIPA buffer (PIERCE, Thermo Scientific) with protease inhibitor cocktail (Roche) and disrupted by sonication on ice. Thirteen microliters of tissue lysates were loaded onto 4-12% NuPAGE gel, transfer onto nitrocellulose membrane and probe for m4 expression using rabbit anti-HA antibody (mAb clone C29F, Cell signaling) and b-actin (mAb clone13E5, cell signaling) for protein loading. Membrane hybridization and detection were performed using PIERCE Fast Western blotting kit Super signal, West pico Rabbit (Thermo Scientific) per manufacturer protocol and imaged using ChemiDoc Imaging system (BIO-RAD).

### Inflammatory cell foci (ICF) quantification

Liver tissues were paraffin-embedded, sectioned and H&E stained by the Experimental histopathology shared resources of the Fred Hutchinson Cancer Center. ICF were counted by a blinded observer and expressed as the number of ICF per surface area which was determined using Fiji.^60^

### Grading of neuronal changes within trigeminal ganglia

Trigeminal ganglia were paraffin-embedded, sectioned, and H&E stained by the Experimental histopathology shared resources of the Fred Hutchinson Cancer Center. Microscopic changes were graded as to severity by a veterinarian pathologist using a standard grading system whereby 0 = no significant change, 1 = minimal, 2 = mild, 3 = moderate, and 4 = severe.

#### Statistical analysis

Statistical analyses for each individual experiment were performed using GraphPad Prism version 9.4.1. Tests were two-sided and p-values smaller than 0.05 considered significant. The specific test used for each analysis is indicated in the corresponding figure legend.

## ACKNOWLEDGMENTS

We are grateful to Cellectis (Paris, France) for the original development of meganuclease m4. This work was supported by National Institutes of Health grant R01AI132599 (KRJ), Caladan Foundation (KRJ), National Institutes of Health grant 5P50AR065139 (Viral Vector Core of the Wellstone Muscular Dystrophy Specialized Research Center, Seattle), National Institutes of Health / National Cancer Institutes, Cancer Center Support Grant P30 CA015704 (Fred Hutchinson Cancer Center Shared Resources Division), donations from the Kenneth Hill Foundation, Krieger Family Trust, Tiny Foundation, and over 2,000 individual donors.

## AUTHOR CONTRIBUTIONS

N.-G.K., M.A., D.S., and K.R.J. conceived this project, N.-G.K., M.A., and K.R.J. wrote the manuscript, N.-G.K. and M.A. designed the experiments, M.A., A.K.H., and P.A.M conducted mouse experiments, H.G.K. and B.E.N., performed NGS, L.S. and L.M.K. performed ddPCR, H.X. and A.C.P supervised the NGS and ddPCR processes. N.-G.K., M.A., M.W., and P.R. analyzed and interpreted data. All authors reviewed and/or contributed to editing the final manuscript.

---

**Table S1.**
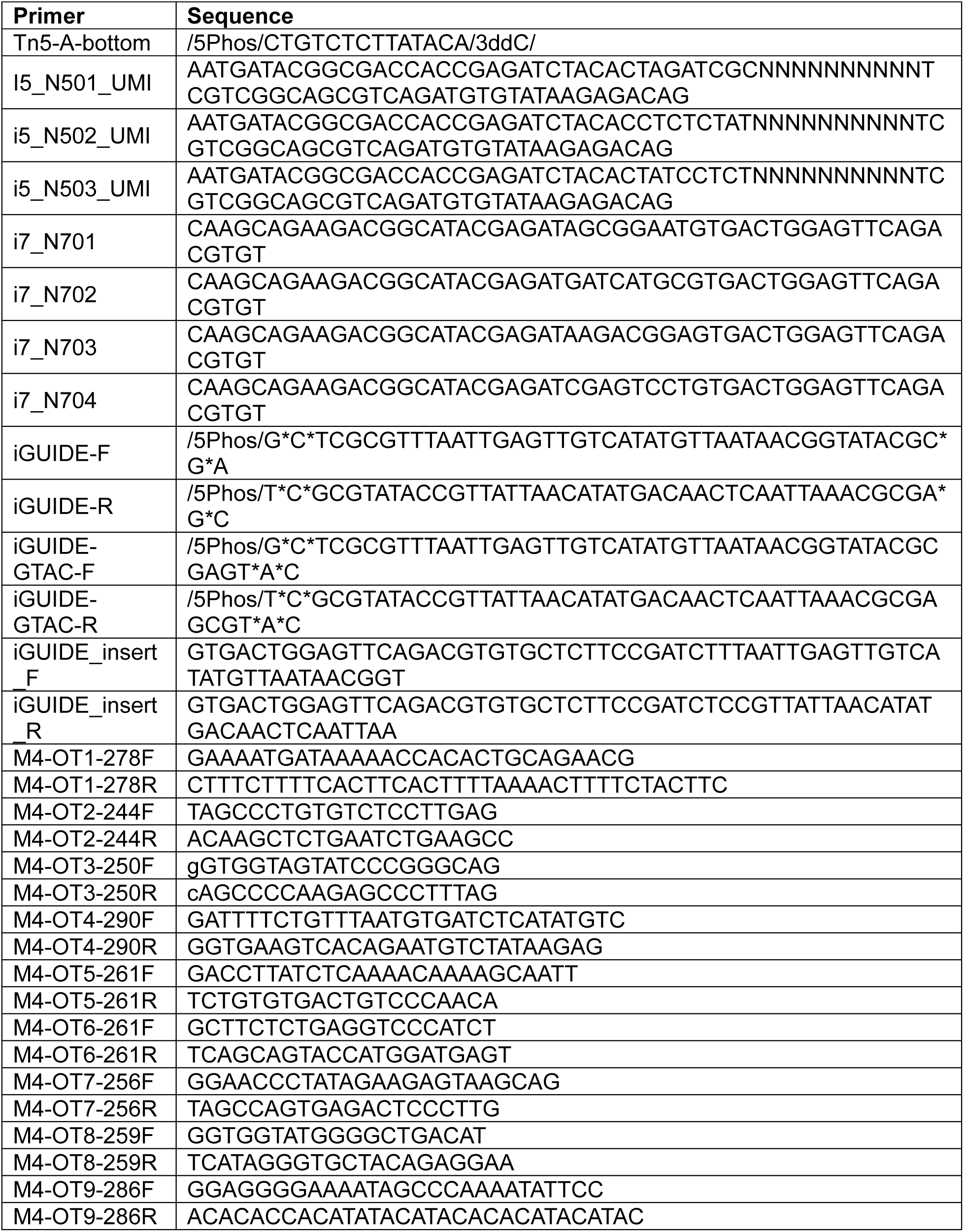
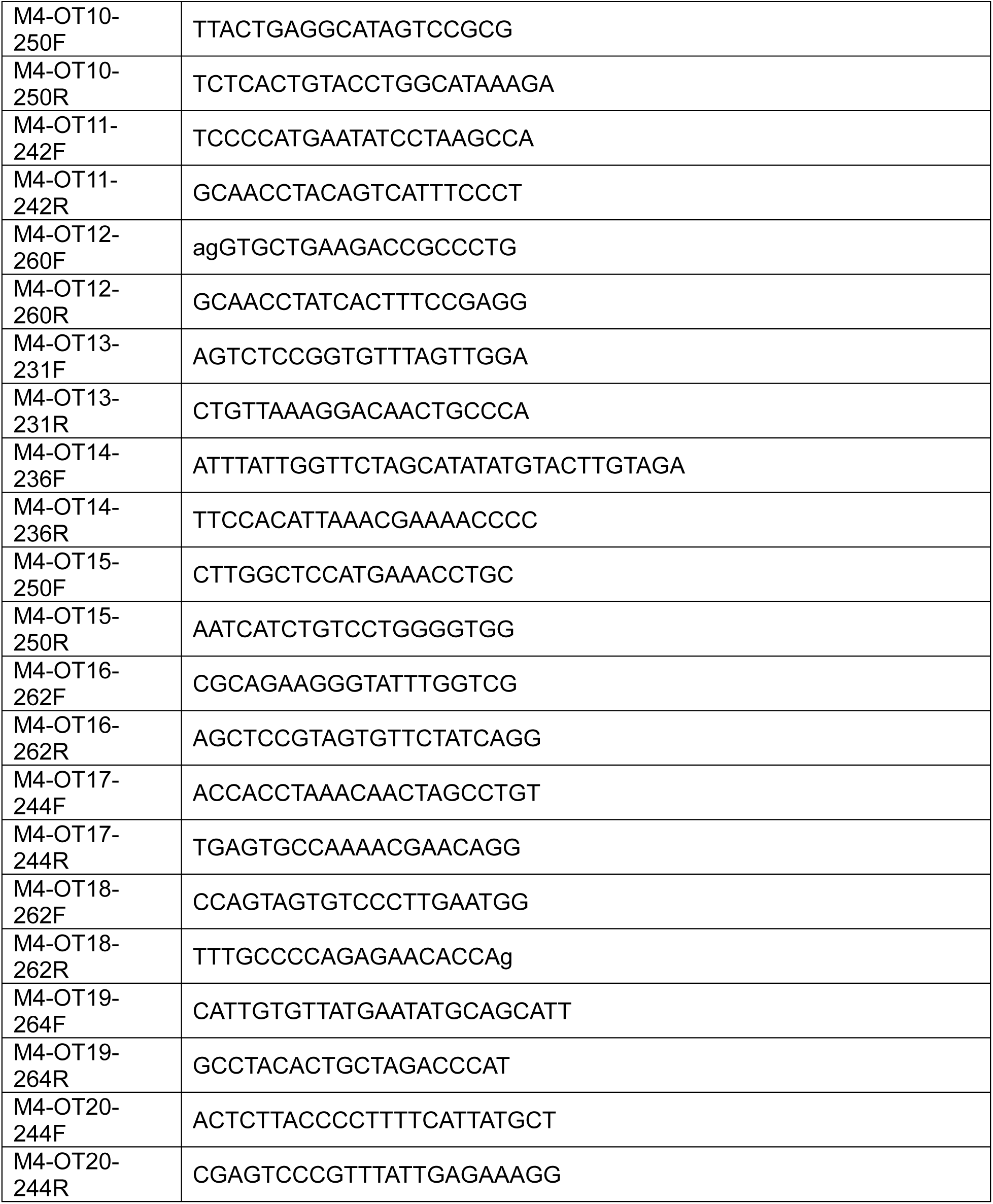
Sequences of primers.

**Figure S1.**
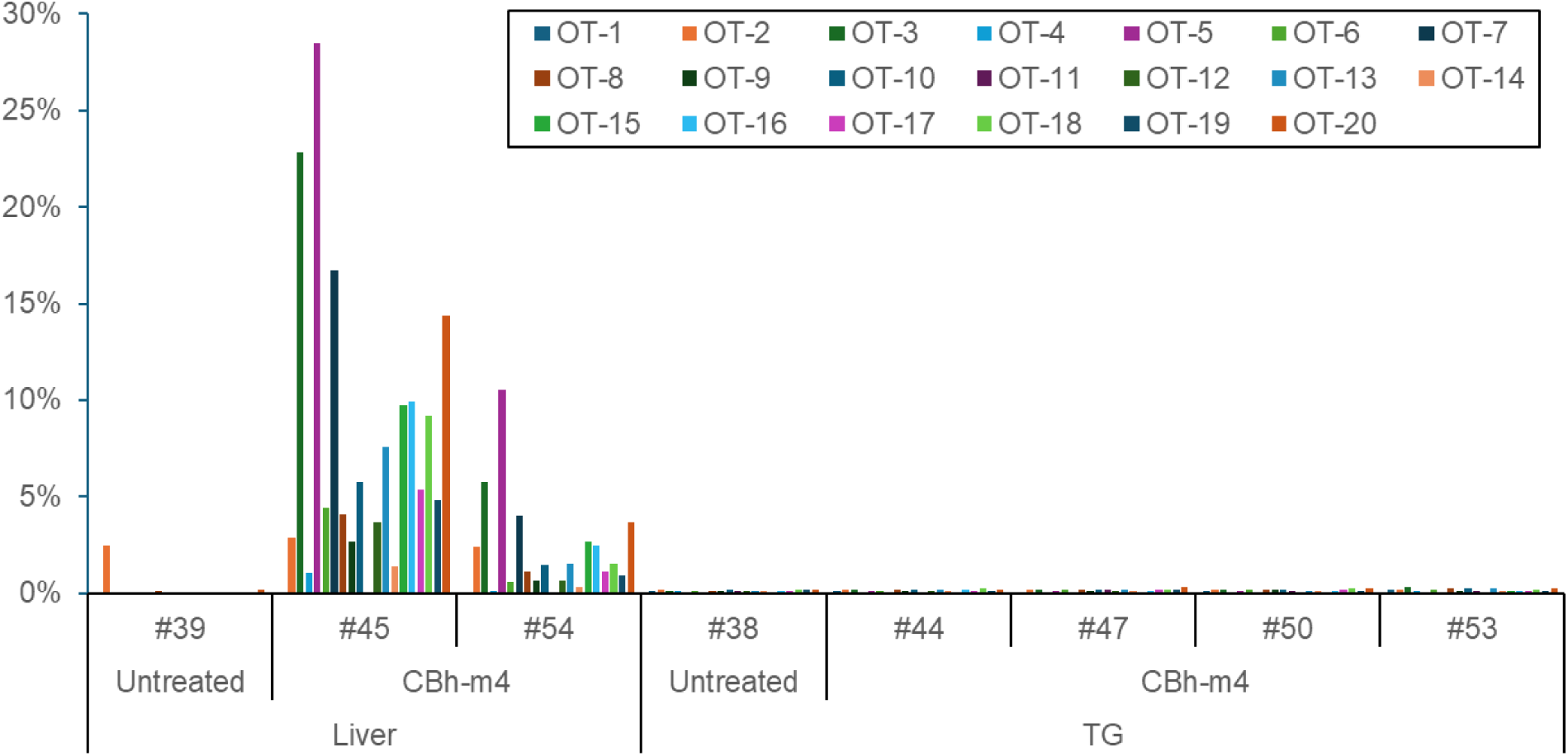
Detection of off-target editing rates in liver or TG tissue DNA using amplicon deep sequencing. Editing rates of the 20 off-target sites identified by GUIDE-Tag were determined through amplicon sequencing of liver (n=2) and TG (n=4) tissue DNA from mice administered AAV9- CBh-m4, which expresses m4 from the ubiquitous CBh promoter.

## REFERENCES

1. Schambach, A., Buchholz, C.J., Torres-Ruiz, R., Cichutek, K., Morgan, M., Trapani, I., and Büning, H. (2024). A new age of precision gene therapy. Lancet Lond. Engl. 403, 568–582. 10.1016/S0140-6736(23)01952-9.

2. Maeder, M.L., and Gersbach, C.A. (2016). Genome-editing Technologies for Gene and Cell Therapy. Mol. Ther. J. Am. Soc. Gene Ther. 24, 430–446. 10.1038/mt.2016.10.

3. Porteus, M.H. (2019). A New Class of Medicines through DNA Editing. N. Engl. J. Med. 380, 947–959. 10.1056/NEJMra1800729.

4. Uddin, F., Rudin, C.M., and Sen, T. (2020). CRISPR Gene Therapy: Applications, Limitations, and Implications for the Future. Front. Oncol. 10, 1387. 10.3389/fonc.2020.01387.

5. Macarrón Palacios, A., Korus, P., Wilkens, B.G.C., Heshmatpour, N., and Patnaik, S.R. (2024). Revolutionizing in vivo therapy with CRISPR/Cas genome editing: breakthroughs, opportunities and challenges. Front. Genome Ed. 6, 1342193. 10.3389/fgeed.2024.1342193.

6. Li, T., Yang, Y., Qi, H., Cui, W., Zhang, L., Fu, X., He, X., Liu, M., Li, P.-F., and Yu, T. (2023). CRISPR/Cas9 therapeutics: progress and prospects. Signal Transduct. Target. Ther. 8, 36. 10.1038/s41392-023-01309-7.

7. Awasthi, R., Maier, H.J., Zhang, J., and Lim, S. (2023). Kymriah® (tisagenlecleucel) - An overview of the clinical development journey of the first approved CAR-T therapy. Hum. Vaccines Immunother. 19, 2210046. 10.1080/21645515.2023.2210046.

8. Frangoul, H., Altshuler, D., Cappellini, M.D., Chen, Y.-S., Domm, J., Eustace, B.K., Foell, J., de la Fuente, J., Grupp, S., Handgretinger, R., et al. (2021). CRISPR-Cas9 Gene Editing for Sickle Cell Disease and β-Thalassemia. N. Engl. J. Med. 384, 252–260. 10.1056/NEJMoa2031054.

9. Kanter, J., Thompson, A.A., Pierciey, F.J., Hsieh, M., Uchida, N., Leboulch, P., Schmidt, M., Bonner, M., Guo, R., Miller, A., et al. (2023). Lovo-cel gene therapy for sickle cell disease: Treatment process evolution and outcomes in the initial groups of the HGB- 206 study. Am. J. Hematol. 98, 11–22. 10.1002/ajh.26741.

10. Raguram, A., Banskota, S., and Liu, D.R. (2022). Therapeutic in vivo delivery of gene editing agents. Cell 185, 2806–2827. 10.1016/j.cell.2022.03.045.

11. Tsuchida, C.A., Wasko, K.M., Hamilton, J.R., and Doudna, J.A. (2024). Targeted nonviral delivery of genome editors in vivo. Proc. Natl. Acad. Sci. U. S. A. 121, e2307796121. 10.1073/pnas.2307796121.

12. Charlesworth, C.T., Hsu, I., Wilkinson, A.C., and Nakauchi, H. (2022). Immunological barriers to haematopoietic stem cell gene therapy. Nat. Rev. Immunol. 22, 719–733. 10.1038/s41577-022-00698-0.

13. Bogdanove, A.J., Bohm, A., Miller, J.C., Morgan, R.D., and Stoddard, B.L. (2018). Engineering altered protein-DNA recognition specificity. Nucleic Acids Res. 46, 4845– 4871. 10.1093/nar/gky289.

14. Laforet, M., McMurrough, T.A., Vu, M., Brown, C.M., Zhang, K., Junop, M.S., Gloor, G.B., and Edgell, D.R. (2019). Modifying a covarying protein-DNA interaction changes substrate preference of a site-specific endonuclease. Nucleic Acids Res. 47, 10830– 10841. 10.1093/nar/gkz866.

15. Zhang, X., Blumenthal, R.M., and Cheng, X. (2024). Updated understanding of the protein-DNA recognition code used by C2H2 zinc finger proteins. Curr. Opin. Struct. Biol. 87, 102836. 10.1016/j.sbi.2024.102836.

16. Boissel, S., Jarjour, J., Astrakhan, A., Adey, A., Gouble, A., Duchateau, P., Shendure, J., Stoddard, B.L., Certo, M.T., Baker, D., et al. (2014). megaTALs: a rare-cleaving nuclease architecture for therapeutic genome engineering. Nucleic Acids Res. 42, 2591–2601. 10.1093/nar/gkt1224.

17. Stone, D., Meumann, N., Kuhlmann, A.-S., Peterson, C.W., Xie, H., Roychoudhury, P., Loprieno, M.A., Vu, X.-K., Strongin, D.E., Kenkel, E.J., et al. (2023). A multiplexed barcode approach to simultaneously evaluate gene delivery by adeno-associated virus capsid variants in nonhuman primates. Hepatol. Commun. 7, e0009. 10.1097/HC9.0000000000000009.

18. Eftekhari, Z., Zohrabi, H., Oghalaie, A., Ebrahimi, T., Shariati, F.S., Behdani, M., and Kazemi-Lomedasht, F. (2024). Advancements and challenges in mRNA and ribonucleoprotein-based therapies: From delivery systems to clinical applications. Mol. Ther. Nucleic Acids 35, 102313. 10.1016/j.omtn.2024.102313.

19. Artemyev, V., Gubaeva, A., Paremskaia, A.I., Dzhioeva, A.A., Deviatkin, A., Feoktistova, S.G., Mityaeva, O., and Volchkov, P.Y. (2024). Synthetic Promoters in Gene Therapy: Design Approaches, Features and Applications. Cells 13, 1963. 10.3390/cells13231963.

20. Guo, Z., Maki, M., Ding, R., Yang, Y., Zhang, B., and Xiong, L. (2014). Genome-wide survey of tissue-specific microRNA and transcription factor regulatory networks in 12 tissues. Sci. Rep. 4, 5150. 10.1038/srep05150.

21. Aubert, M., Haick, A.K., Strongin, D.E., Klouser, L.M., Loprieno, M.A., Stensland, L., Santo, T.K., Huang, M.-L., Hyrien, O., Stone, D., et al. (2024). Gene editing for latent herpes simplex virus infection reduces viral load and shedding in vivo. Nat. Commun. 15, 4018. 10.1038/s41467-024-47940-y.

22. Aubert, M., Strongin, D.E., Roychoudhury, P., Loprieno, M.A., Haick, A.K., Klouser, L.M., Stensland, L., Huang, M.-L., Makhsous, N., Tait, A., et al. (2020). Gene editing and elimination of latent herpes simplex virus in vivo. Nat. Commun. 11, 4148. 10.1038/s41467-020-17936-5.

23. Ertl, H.C.J. (2022). Immunogenicity and toxicity of AAV gene therapy. Front. Immunol. 13, 975803. 10.3389/fimmu.2022.975803.

24. Liang, S.-Q., Liu, P., Smith, J.L., Mintzer, E., Maitland, S., Dong, X., Yang, Q., Lee, J., Haynes, C.M., Zhu, L.J., et al. (2022). Genome-wide detection of CRISPR editing in vivo using GUIDE-tag. Nat. Commun. 13, 437. 10.1038/s41467-022-28135-9.

25. Tsai, S.Q., Zheng, Z., Nguyen, N.T., Liebers, M., Topkar, V.V., Thapar, V., Wyvekens, N., Khayter, C., Iafrate, A.J., Le, L.P., et al. (2015). GUIDE-seq enables genome-wide profiling of off-target cleavage by CRISPR-Cas nucleases. Nat. Biotechnol. 33, 187–197. 10.1038/nbt.3117.

26. Musunuru, K., Chadwick, A.C., Mizoguchi, T., Garcia, S.P., DeNizio, J.E., Reiss, C.W., Wang, K., Iyer, S., Dutta, C., Clendaniel, V., et al. (2021). In vivo CRISPR base editing of PCSK9 durably lowers cholesterol in primates. Nature 593, 429–434. 10.1038/s41586-021-03534-y.

27. Wienert, B., Wyman, S.K., Richardson, C.D., Yeh, C.D., Akcakaya, P., Porritt, M.J., Morlock, M., Vu, J.T., Kazane, K.R., Watry, H.L., et al. (2019). Unbiased detection of CRISPR off-targets in vivo using DISCOVER-Seq. Science 364, 286–289. 10.1126/science.aav9023.

28. Nobles, C.L., Reddy, S., Salas-McKee, J., Liu, X., June, C.H., Melenhorst, J.J., Davis, M.M., Zhao, Y., and Bushman, F.D. (2019). iGUIDE: an improved pipeline for analyzing CRISPR cleavage specificity. Genome Biol. 20, 14. 10.1186/s13059-019-1625-3.

29. Grosse, S., Huot, N., Mahiet, C., Arnould, S., Barradeau, S., Clerre, D.L., Chion-Sotinel, I., Jacqmarcq, C., Chapellier, B., Ergani, A., et al. (2011). Meganuclease-mediated Inhibition of HSV1 Infection in Cultured Cells. Mol. Ther. J. Am. Soc. Gene Ther. 19, 694– 702. 10.1038/mt.2010.302.

30. Arnould, S., Delenda, C., Grizot, S., Desseaux, C., Pâques, F., Silva, G.H., and Smith, J. (2011). The I-CreI meganuclease and its engineered derivatives: applications from cell modification to gene therapy. Protein Eng. Des. Sel. PEDS 24, 27–31. 10.1093/protein/gzq083.

31. Liu, Y., Zou, R.S., He, S., Nihongaki, Y., Li, X., Razavi, S., Wu, B., and Ha, T. (2020). Very fast CRISPR on demand. Science 368, 1265–1269. 10.1126/science.aay8204.

32. Gray, S.J., Foti, S.B., Schwartz, J.W., Bachaboina, L., Taylor-Blake, B., Coleman, J., Ehlers, M.D., Zylka, M.J., McCown, T.J., and Samulski, R.J. (2011). Optimizing promoters for recombinant adeno-associated virus-mediated gene expression in the peripheral and central nervous system using self-complementary vectors. Hum. Gene Ther. 22, 1143–1153. 10.1089/hum.2010.245.

33. Hioki, H., Kameda, H., Nakamura, H., Okunomiya, T., Ohira, K., Nakamura, K., Kuroda, M., Furuta, T., and Kaneko, T. (2007). Efficient gene transduction of neurons by lentivirus with enhanced neuron-specific promoters. Gene Ther. 14, 872–882. 10.1038/sj.gt.3302924.

34. Salk, J.J., Schmitt, M.W., and Loeb, L.A. (2018). Enhancing the accuracy of next-generation sequencing for detecting rare and subclonal mutations. Nat. Rev. Genet. 19, 269–285. 10.1038/nrg.2017.117.

35. Brown, B.D., and Naldini, L. (2009). Exploiting and antagonizing microRNA regulation for therapeutic and experimental applications. Nat. Rev. Genet. 10, 578–585. 10.1038/nrg2628.

36. Linsen, S.E., Tops, B.B., and Cuppen, E. (2008). miRNAs: small changes, widespread effects. Cell Res. 18, 1157–1159. 10.1038/cr.2008.311.

37. Geisler, A., Jungmann, A., Kurreck, J., Poller, W., Katus, H.A., Vetter, R., Fechner, H., and Müller, O.J. (2011). microRNA122-regulated transgene expression increases specificity of cardiac gene transfer upon intravenous delivery of AAV9 vectors. Gene Ther. 18, 199–209. 10.1038/gt.2010.141.

38. Jopling, C. (2012). Liver-specific microRNA-122: Biogenesis and function. RNA Biol. 9, 137–142. 10.4161/rna.18827.

39. Pacesa, M., Pelea, O., and Jinek, M. (2024). Past, present, and future of CRISPR genome editing technologies. Cell 187, 1076–1100. 10.1016/j.cell.2024.01.042.

40. Gillmore, J.D., Gane, E., Taubel, J., Kao, J., Fontana, M., Maitland, M.L., Seitzer, J., O’Connell, D., Walsh, K.R., Wood, K., et al. (2021). CRISPR-Cas9 In Vivo Gene Editing for Transthyretin Amyloidosis. N. Engl. J. Med. 385, 493–502. 10.1056/NEJMoa2107454.

41. Longhurst, H.J., Lindsay, K., Petersen, R.S., Fijen, L.M., Gurugama, P., Maag, D., Butler, J.S., Shah, M.Y., Golden, A., Xu, Y., et al. (2024). CRISPR-Cas9 In Vivo Gene Editing of KLKB1 for Hereditary Angioedema. N. Engl. J. Med. 390, 432–441. 10.1056/NEJMoa2309149.

42. Lee, R.G., Mazzola, A.M., Braun, M.C., Platt, C., Vafai, S.B., Kathiresan, S., Rohde, E., Bellinger, A.M., and Khera, A.V. (2023). Efficacy and Safety of an Investigational Single-Course CRISPR Base-Editing Therapy Targeting PCSK9 in Nonhuman Primate and Mouse Models. Circulation 147, 242–253. 10.1161/CIRCULATIONAHA.122.062132.

43. Kitawi, R., Ledger, S., Kelleher, A.D., and Ahlenstiel, C.L. (2024). Advances in HIV Gene Therapy. Int. J. Mol. Sci. 25, 2771. 10.3390/ijms25052771.

44. Di, J., Du, Z., Wu, K., Jin, S., Wang, X., Li, T., and Xu, Y. (2022). Biodistribution and Non-linear Gene Expression of mRNA LNPs Affected by Delivery Route and Particle Size. Pharm. Res. 39, 105–114. 10.1007/s11095-022-03166-5.

45. Simoni, C., Barbon, E., Muro, A.F., and Cantore, A. (2024). In vivo liver targeted genome editing as therapeutic approach: progresses and challenges. Front. Genome Ed. 6, 1458037. 10.3389/fgeed.2024.1458037.

46. Berta, T., Strong, J.A., Zhang, J.-M., and Ji, R.-R. (2023). Targeting dorsal root ganglia and primary sensory neurons for the treatment of chronic pain: an update. Expert Opin. Ther. Targets 27, 665–678. 10.1080/14728222.2023.2247563.

47. Yang, G., Si-Tayeb, K., Corbineau, S., Vernet, R., Gayon, R., Dianat, N., Martinet, C., Clay, D., Goulinet-Mainot, S., Tachdjian, G., et al. (2013). Integration-deficient lentivectors: an effective strategy to purify and differentiate human embryonic stem cell-derived hepatic progenitors. BMC Biol. 11, 86. 10.1186/1741-7007-11-86.

48. Magnuson, M.A., and Osipovich, A.B. (2013). Pancreas-specific Cre driver lines and considerations for their prudent use. Cell Metab. 18, 9–20. 10.1016/j.cmet.2013.06.011.

49. Bilal, A.S., Thuerauf, D.J., Blackwood, E.A., and Glembotski, C.C. (2022). Design and Production of Heart Chamber-Specific AAV9 Vectors. Methods Mol. Biol. Clifton NJ 2573, 89–113. 10.1007/978-1-0716-2707-5_8.

50. Senturk, S., Shirole, N.H., Nowak, D.G., Corbo, V., Pal, D., Vaughan, A., Tuveson, D.A., Trotman, L.C., Kinney, J.B., and Sordella, R. (2017). Rapid and tunable method to temporally control gene editing based on conditional Cas9 stabilization. Nat. Commun. 8, 14370. 10.1038/ncomms14370.

51. Nakahara, E., Mullapudi, V., Collier, G.E., Joachimiak, L.A., and Hulleman, J.D. (2022). Development of a New DHFR-Based Destabilizing Domain with Enhanced Basal Turnover and Applicability in Mammalian Systems. ACS Chem. Biol. 17, 2877–2889. 10.1021/acschembio.2c00518.

52. Miyazaki, Y., Imoto, H., Chen, L., and Wandless, T.J. (2012). Destabilizing domains derived from the human estrogen receptor. J. Am. Chem. Soc. 134, 3942–3945. 10.1021/ja209933r.

53. Shrivastava, S., Ray, R.M., Holguin, L., Echavarria, L., Grepo, N., Scott, T.A., Burnett, J., and Morris, K.V. (2021). Exosome-mediated stable epigenetic repression of HIV-1. Nat. Commun. 12, 5541. 10.1038/s41467-021-25839-2.

54. McCutcheon, S.R., Rohm, D., Iglesias, N., and Gersbach, C.A. (2024). Epigenome editing technologies for discovery and medicine. Nat. Biotechnol. 42, 1199–1217. 10.1038/s41587-024-02320-1.

55. Research, C. for B.E. and (2024). Human Gene Therapy Products Incorporating Human Genome Editing. https://www.fda.gov/regulatory-information/search-fda-guidance-documents/human-gene-therapy-products-incorporating-human-genome-editing.

56. Choi, V.W., Asokan, A., Haberman, R.A., and Samulski, R.J. (2007). Production of recombinant adeno-associated viral vectors for in vitro and in vivo use. Curr. Protoc. Mol. Biol. Chapter 16, Unit 16.25. 10.1002/0471142727.mb1625s78.

57. Zolotukhin, S., Byrne, B.J., Mason, E., Zolotukhin, I., Potter, M., Chesnut, K., Summerford, C., Samulski, R.J., and Muzyczka, N. (1999). Recombinant adeno-associated virus purification using novel methods improves infectious titer and yield. Gene Ther. 6, 973–985. 10.1038/sj.gt.3300938.

58. Aurnhammer, C., Haase, M., Muether, N., Hausl, M., Rauschhuber, C., Huber, I., Nitschko, H., Busch, U., Sing, A., Ehrhardt, A., et al. (2012). Universal real-time PCR for the detection and quantification of adeno-associated virus serotype 2-derived inverted terminal repeat sequences. Hum. Gene Ther. Methods 23, 18–28. 10.1089/hgtb.2011.034.

59. Aubert, M., Madden, E.A., Loprieno, M., DeSilva Feelixge, H.S., Stensland, L., Huang, M.-L., Greninger, A.L., Roychoudhury, P., Niyonzima, N., Nguyen, T., et al. (2016). In vivo disruption of latent HSV by designer endonuclease therapy. JCI Insight 1, e88468. 10.1172/jci.insight.88468.

60. Schindelin, J., Arganda-Carreras, I., Frise, E., Kaynig, V., Longair, M., Pietzsch, T., Preibisch, S., Rueden, C., Saalfeld, S., Schmid, B., et al. (2012). Fiji: an open-source platform for biological-image analysis. Nat. Methods 9, 676–682. 10.1038/nmeth.2019.

